# ProCyon: A multimodal foundation model for protein phenotypes

**DOI:** 10.1101/2024.12.10.627665

**Authors:** Owen Queen, Yepeng Huang, Robert Calef, Valentina Giunchiglia, Tianlong Chen, George Dasoulas, LeAnn Tai, Gianmarco Abbadessa, Owain Howell, Michelle M. Li, Yasha Ektefaie, Ayush Noori, Ildiko Farkas, Joseph Brown, Tom Cobley, Karin Hrovatin, Tom Hartvigsen, Fabian J. Theis, Bradley L. Pentelute, James Zou, Vikram Khurana, David Owen, Richard Nicholas, Manolis Kellis, Marinka Zitnik

## Abstract

Characterizing human proteins remains a major challenge: approximately 29% of human proteins lack experimentally validated functions and even well-annotated proteins often lack context-specific phenotypic insights. To enable universal modeling of protein phenotypes, we present ProCyon, a multimodal foundation model that utilizes protein sequence, structure, and natural language for generating and predicting protein phenotypes across diverse knowledge domains. ProCyon is trained on our novel dataset, ProCyon-Instruct, with 33 million protein phenotype instructions. On dozens of benchmarking tasks, ProCyon performs competitively against single-modal and multimodal models. Further, ProCyon conditionally retrieves proteins via mechanisms of action of small molecule drugs and disease contexts, and it generates candidate phenotypic descriptions for poorly characterized proteins, including those implicated in Parkinson’s disease that were identified after ProCyon’s knowledge cutoff date. We experimentally confirm ProCyon’s predictions in multiple sclerosis using post-mortem brain RNA-seq, identifying novel MS genes and elucidating associated pathway mechanisms consistent with cortical pathology. ProCyon paves the way toward a general approach to generate functional insights into the human proteome.

---

Decoding protein function remains a fundamental challenge for understanding how living systems operate at the molecular level. Despite advances in characterizing protein sequences and predicting protein structures [1], approximately 7% of human protein-coding genes still lack proteomic evidence [2, 3], and even well-annotated proteins often lack context-specific phenotypic insights. The 2024 HUPO Human Proteome Project (HPP) reports 5,656 proteins (29%) without experimentally validated functions in the FE4 and FE5 tiers [3], while 95% of life science publications focus on only 5,000 proteins [4].

Protein phenotypes—observable properties and characteristics resulting from a protein’s function—are multiscale and encompass molecular processes, therapeutic interactions, cellular pathways, organismal traits, and diseases [5–7]. Since these context-dependent phenotypes can vary under different biological conditions, characterizing proteins is a challenging endeavor [8–10]. Although valuable [11, 12], existing approaches for predicting protein function often do not capture the full spectrum of protein phenotypes or generate overly generic annotations (e.g., ATP binding, nucleic acid binding) [13]. Addressing this challenge is a central focus of initiatives such as the HPP Grand Challenge, “A Function for Each Protein” [13], which aims to determine phenotypes of every protein in the context of human health and disease.

Traditionally, protein function prediction methods rely on transferring functional knowledge between homologs identified via sequence or structure similarity. Although homology captures evolutionary conservation, homologs can exhibit different functions in distinct tissues and organisms [14]. While recent work, such as protein language models [15,16] and representation learning methods [17–19], generate embeddings that better capture functional similarity across proteins, the implicit reliance on homology alone remains. Additionally, sequence- and structure-based methods rely on pre-defined functional categories for prediction, limiting their ability to generalize to phenotypes not yet codified in controlled ontologies and pre-defined vocabularies of protein function. These narrow definitions of protein function also render the models unable to capture proteins’ multiscale, context-dependent phenotypes, which span knowledge domains like molecular function, disease associations, and molecular interactions [6, 20–22].

Leveraging natural language descriptions presents a promising avenue for flexibly modeling protein phenotypes and enabling human-AI collaboration. However, even advanced large language models (LLMs) [23] learn only from text-based associations, typically derived from biomedical literature, represent proteins through static identifiers such as UniProt IDs [24] or HGNC gene sym-bols [25], and cannot incorporate non-textual protein data, such as atom-level protein structures. These factors restrict LLMs’ ability to generalize to new proteins [26] and bias them against poorly characterized proteins [27]. Some models [28–30] have explored jointly learning on proteins and natural language, but they remain limited in scope by the diversity of phenotypes they are trained on and have limited text generation capabilities.

Here, we introduce ProCyon, a multimodal foundation model for contextual protein retrieval and generation of protein phenotype descriptions. ProCyon leverages natural language as the unifying modality to bridge knowledge domains and represent arbitrary protein phenotypes. ProCyon models phenotypes using inputs containing interleaved textual phenotype descriptions, protein sequences, protein structures, and small molecular structures. ProCyon is not constrained by controlled vocabularies or pre-defined terminologies of protein function, and can generate open-ended phenotype descriptions beyond those seen during training. To train ProCyon, we have developed ProCyon-INSTRUCT, an instruction tuning dataset consisting of over 33 million examples linking proteins to diverse phenotypes. By learning a unified latent space of proteins and phenotypes, ProCyon can flexibly adapt to a wide range of tasks previously achievable only by specialized models, including tasks that require connecting phenotypes across scales, from molecular characteristics to organism-wide traits (Figure 1a).

**Figure 1.**
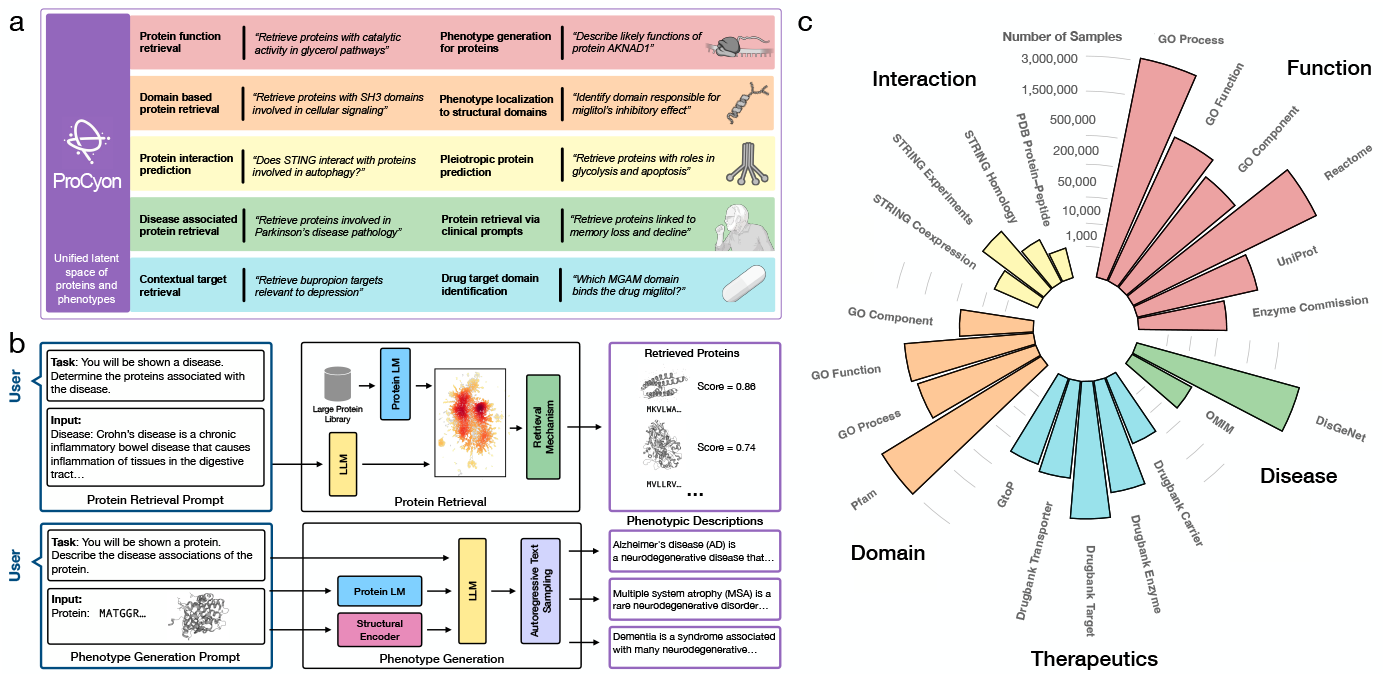
Overview of ProCyon model architecture and ProCyon-Instruct dataset. **a)** ProCyon models proteins and phenotypes in a unified latent space, enabling wide applications across knowledge domains (shown are example prompts). **b)** ProCyon is a multimodal foundation model consisting of a large language model and multimodal molecular encoders, trained to perform both protein retrieval and phenotype generation. For protein retrieval, a prompt is processed into the LLM and compared against a library of proteins. ProCyon then outputs a ranked list of proteins, domains, peptides, or polypeptides. For phenotype generation, the input text and multimodal elements are processed by the LLM via multimodal token decomposition. ProCyon then performs autoregressive text generation to generate protein phenotypes and other free-form answers. **c)** A comprehensive ProCyon-INSTRUCT dataset with 33,899,528 protein-phenotype instructions is curated from five knowledge domains (Function, Therapeutics, Disease, Protein Domains, and Interaction) and used to train ProCyon. Details of dataset curation, phenotype description rephrasing, and instruction templating can be found in Extended Data Fig. 4, Methods Sec. 1 and 3.1, and Supplementary Note 2. The number displayed shows the number of protein-phenotype description pairs curated from each database.

ProCyon outperforms existing models, including state-of-the-art protein representation and protein-text models, in contextual protein retrieval and question answering on 12 benchmarking tasks spanning multiple knowledge domains. We further demonstrate that ProCyon can generalize to a diverse range of tasks via its natural language interface. ProCyon performs contextual protein retrieval via natural language prompts that condition on different contexts, enabling the retrieval of proteins with complex, pleiotropic phenotypes and the prioritization of disease-associated proteins with increasingly precise disease descriptions. ProCyon is also able to identify drug-binding domains within proteins and predict peptide binders for the ACE2 enzyme tested against an experimental dataset.

We use experimental data to validate ProCyon’s predicted and generated phenotypes for poorly characterized proteins. We find that ProCyon’s predicted phenotypes have strong concordance with annotations derived from perturb-seq data. We show that human expert evaluations support ProCyon’s predicted phenotypes for poorly characterized proteins associated with Parkinson’s disease (PD). Finally, we validate ProCyon’s conditional protein retrieval for multiple sclerosis-associated phenotypes (MS) via RNA sequencing of post-mortem brain samples from MS brain donors. Our training datasets, training and inference code, and pretrained models are publicly available.

## Results

### ProCyon model architecture

ProCyon is a multimodal protein foundation model with 11 billion parameters. Composed of a Llama-3-8B language model [31] and encoders for protein sequence (ESM-2 3B [32]) and structure (GearNet [19]), ProCyon can process inputs of natural language and protein sequence and structure. To support a wide range of downstream tasks, ProCyon has two output modes: protein retrieval and phenotype generation (Figure 1b).

In protein retrieval, ProCyon retrieves a ranked list of proteins that match a user’s prompt. First, the protein language model processes protein sequences and projects them into the unified latent space via a retrieval-specific projector. The natural language prompt is then processed by the LLM and projected into the unified latent space by a query connector, allowing ProCyon to identify the proteins most closely related to the projected prompt (Supplementary Note 1, Extended Data Figure 1).

In phenotype generation, ProCyon generates natural language descriptions of protein phenotypes conditioned on the user’s prompt. The prompt is split into multimodal components: the text is tokenized via a standard text tokenizer, and the protein sequence and structure are encoded via the modality-specific encoders. These multimodal tokens form a unified prompt that is then processed by the LLM, enabling phenotype generation via standard language model sampling techniques. We also leverage ProCyon’s text generation capability for question-answering (QA), where the model is asked whether the phenotype and protein are associated, prompting a “yes” or “no” response. These QA scores are also used as a filtering mechanism (Methods Sec. 2.3, Extended Data Figure 2) to improve phenotype generation (Extended Data Figure 3).

### ProCyon-INSTRUCT dataset with 33 million protein-phenotype instructions

We construct ProCyon-INSTRUCT, an instruction tuning dataset that spans 12 data sources and 5 knowledge domains: (1) **function**, covering molecular and cellular phenotypes; (2) **disease**, covering gene-disease associations; (3) **therapeutics**, covering protein-drug associations; (4) **protein domains**, covering evolutionarily-conserved functional sub-units of proteins; and (5) **interaction**, covering protein-protein and protein-peptide interactions (Methods Sec. 1, Supplementary Note 2, Supplementary Table 1). To increase the diversity of phenotypes, we rephrase the original descriptions using OpenAI’s GPT models. Our augmentation strategy accounts for different levels of description (e.g., rephrasing, summarization, and simplification) as well as user expertise (Supplementary Notes 3-4). This brings our number of protein-phenotype description pairs to 11,347,986 (Figure 1c, Methods Sec. 1.3, Supplementary Tables 2-3).

We transform the protein-phenotype pairs of ProCyon-INSTRUCT into a multimodal instruction tuning dataset. For each protein-phenotype pair, we create an instruction tuning example that consists of a natural language task definition, associated protein inputs, and an expected answer [33–37] (Extended Data Figures 4a and 5). Instructions are constructed for retrieval, QA, and phenotype generation tasks. Retrieval instructions are used to train ProCyon’s retrieval module via contrastive learning (Methods Sec. 3.4). QA and phenotype generation instructions are used for causal language model training [33]. Our final ProCyon-INSTRUCT dataset consists of 33,899,528 multimodal instruction examples (Supplementary Note 5).

Unless otherwise specified, ProCyon refers to the model trained on all data; when used generically, it refers to the overall modeling framework (Supplementary Note 8). We create data splits of ProCyon-INSTRUCT that enable rigorous assessment of ProCyon’s ability to model protein phenotypes. We partition ProCyon-INSTRUCT into three subsets, which we refer to as “many-shot”, “few-shot”, and “zero-shot” phenotypes. We minimize overlap between these subsets by leveraging the hierarchical or networked structure of phenotypes within each data source, ensuring that ontologically similar phenotypes are assigned to the same subset (Methods Sec. 1, Supplementary Note 6). To further prevent information leakage, we use text embeddings of phenotype descriptions to ensure that semantically similar phenotypes are grouped together (Supplementary Note 7, Supplementary Figures 1 and 2). Protein-phenotype pairs are assigned to train or test sets based on the corresponding phenotypes’ shot-level assignments: pairs from many-shot phenotypes are split randomly between train and test; a small number (2 or 5 depending on data source) of pairs from each few-shot phenotype are assigned to the train set and the remainder to the test set; and all pairs from zero-shot phenotypes are assigned to the test set. To assess zero-shot generalization, we train a version of the model, ProCyon-SPLIT, using only the train set and evaluate it on the test set.

### ProCyon accurately retrieves proteins from flexible prompts of phenotypes

To assess ProCyon’s retrieval capabilities (Figure 2a), we construct 6 retrieval benchmarks from the test split of ProCyon-INSTRUCT. Each task entails retrieving proteins based on prompts describing phenotypes. These benchmarks span 6,153 unique phenotypes with 57,818 protein-phenotype pairs. We compare against homology-based annotation methods (BLAST-kNN) [38], deep protein representation learning methods (ESM2-kNN, ESM2-MLP, ESM3-kNN, ESM3-MLP, GearNet-kNN, GearNet-MLP) [16, 19], and multimodal models that integrate protein representations and free-text descriptions (BioTranslator, ProtST) [28, 29] (Supplementary Notes 9-10). ProCyon-SPLIT is the top performing model in 5 out of 6 tasks, and is among the best three models in all tasks (Figure 2b, Supplementary Table 4, Supplementary Figure 3). ProCyon-Split has the highest performance across tasks, with an average *F*_max_ = 0.723 compared to the next-highest *F*_max_ = 0.615 by ESM3-MLP. Its retrieval performance generally decreases on rare (few-shot) or unseen (zero-shot) phenotypes, but remains strong even for unseen phenotypes, with average *F*_max_ values of 0.728, 0.712, and 0.702 on many-, few-, and zero-shot splits, respectively (Figure 2c, Supplementary Table 5).

**Figure 2.**
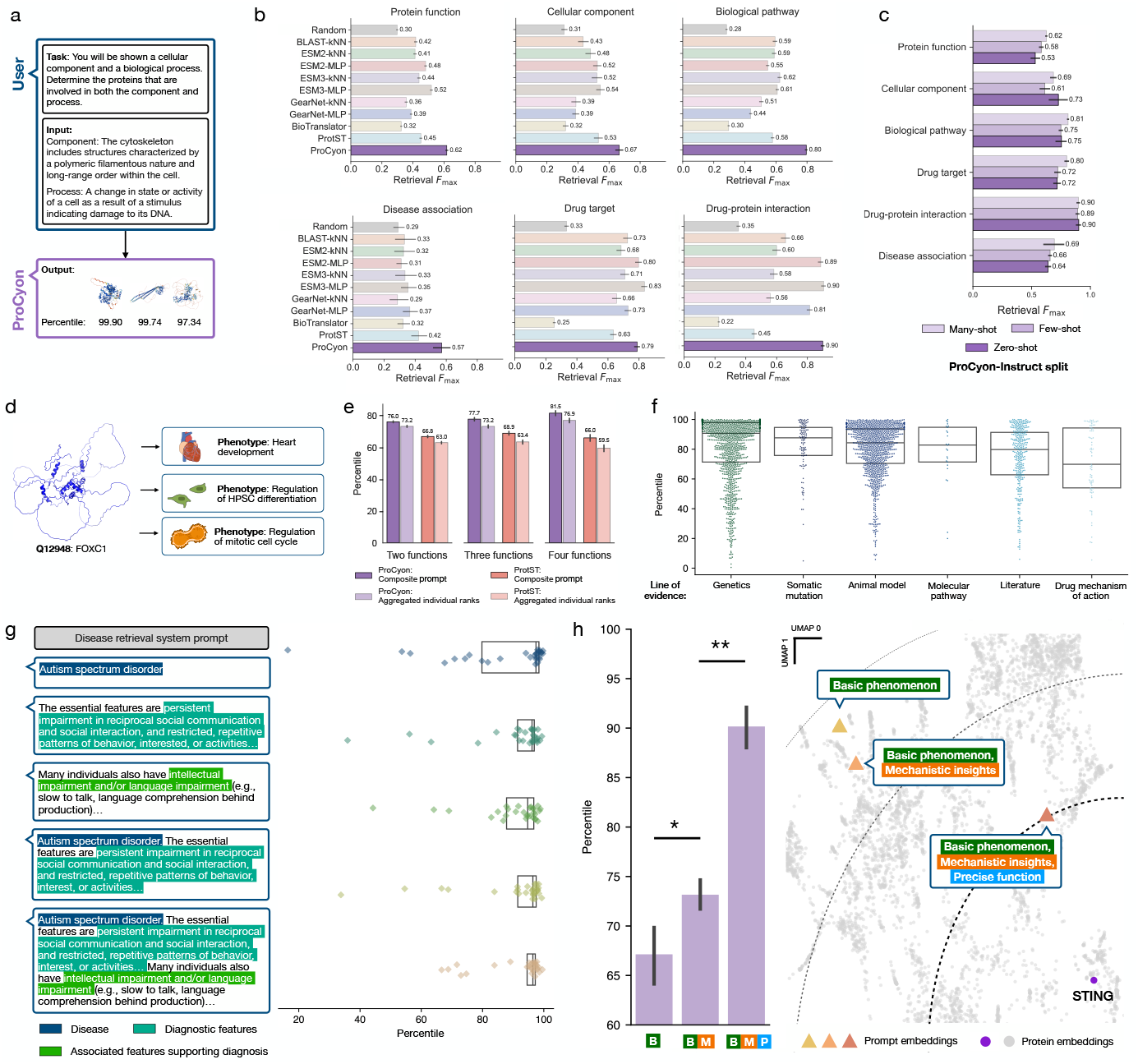
ProCyon accurately retrieves proteins from flexible phenotypes. **a)** In protein retrieval, ProCyon receives a phenotype as input and outputs a ranked list of proteins, domains, peptides, or polypeptides. **b)** Benchmarking ProCyon’s retrieval capabilities against other approaches across distinct knowledge domains. *x* axes show *F*_max_, the maximum *F*_1_ score across any cutoff. Error bars show bootstrapped 95% confidence intervals. One retrieval task corresponds to retrieving proteins for one phenotype. **c)** ProCyon’s retrieval performance across splits of ProCyon-INSTRUCT. **d)** Illustration of pleiotropic protein functions. The FOXC1 protein is implicated in heart development, regulation of HPSC differentiation, and regulation of the mitotic cell cycle. **e)** Percentiles of potentially pleiotropic proteins ranked by ProCyon using composite prompts composed of two, three, and four pathways, in comparison with aggregated individual ranks. Also compared with ProtST. All comparisons have *p*-value *<* 0.001; two-sided Wilcoxon signed-rank test. **f)** External validation of disease-associated protein retrieval across different lines of evidence for association. *y* axis shows median retrieval percentile scores for each disease across different lines of evidence. **g)** Protein retrieval with different disease descriptions. Left, example texts for prompts with gradually increasing information (Disease, Diagnostics, Associated Features supporting Diagnosis and their combinations) for Autistic Spectrum Disorder. Right, median retrieval percentile scores derived for each disease with increasingly informative prompts. **h)** Left, retrieval percentile of STING as more precise prompts are provided as input. Right, UMAP of unified latent space of phenotypes and proteins, highlighting prompt embeddings and STING protein embedding. Dashed contours indicate distance from STING protein embedding; thicker lines denote closer distance. * *p*-value *<* 0.05, ** *p*-value *<* 0.005; two-sided Mann-Whitney U test.

Next, we evaluate ProCyon’s retrieval of pleiotropic proteins involved in multiple biological pathways (Methods Sec. 4.5). Proteins can have multiple functions or phenotypes, a phenomenon known as pleiotropy [39]. One such protein is FOXC1, which is involved in heart development, regulation of hematopoietic stem and progenitor cell (HSPC) differentiation, and regulation of the mitotic cell cycle (Figure 2d). We design a pleiotropic retrieval task where the model is prompted with descriptions of multiple phenotypes (up to four) and asked to prioritize proteins involved in all phenotypes (“composite prompt”). ProCyon is never explicitly trained on pleiotropic function retrieval. We compare composite prompts to aggregating retrieved lists from individual phenotype prompts (“aggregated individual ranks”) and to ProtST on the same task using both aggregation methods. In all settings (Figure 2e), ProCyon has more accurate retrieval than other methods (*p*-value *<* 10^*−*4^; Wilcoxon signed rank test), and the composite prompt outperforms aggregated individual ranks (*p*-value *<* 10^*−*4^; Wilcoxon signed rank test). ProCyon also retrieves proteins involved with crosstalk between signaling pathways [40], again with the composite prompt outperforming the aggregated individual ranks (*p*-value = 0.026; Wilcoxon signed rank test) (Supplementary Note 11, Supplementary Figure 4; Methods Sec. 4.5).

To further probe retrieval of disease-associated proteins, we stratify performance by lines of evidence linking the protein to the disease (Supplementary Note 12). ProCyon-SPLIT highly ranks disease-associated proteins across all evidence types, as measured by high average retrieval percentiles from the library of 18,174 human proteins (Figure 2f).

We also measure ProCyon’s ability to leverage increasingly detailed prompts to retrieve disease-associated proteins (Figure 2g). Using descriptions from the Diagnostic and Statistical Manual of Mental Disorders (DSM-5) [41], we incrementally expand prompts from the disease name alone (“Disease”) to include “Diagnostic Features” and “Associated Features Supporting Diagnosis.” ProCyon-SPLIT has improved retrieval performance as the level of detail in the prompt increases, reaching up to 91.69*±*10.15 retrieval percentile when all information is provided, with a significant reduction in variance (*p* = 0.01, Bartlett’s test when comparing across levels of details; Supplementary Figure 5). Similarly, ProCyon-SPLIT retains high performance on 22 monogenic diseases even when excluding disease name from the description (Extended Data Figure 6, Supplementary Table 6), suggesting that ProCyon captures relationships between proteins and disease phenotypes beyond simple name matching.

Finally, we challenge ProCyon with prompts of phenotypes that do not correspond to protein functions codified in controlled vocabularies or pre-defined ontologies. We first demonstrate that ProCyon successfully retrieves proteins associated with distinct cognitive functions—critical phenotypic features not captured by existing gene-disease association databases (Supplementary Note 13, Extended Data Figure 7). Next, we prompt ProCyon with descriptions about neuronal inflammatory stress response (NISR) to evaluate whether it can prioritize the STING1 protein, whose role in NISR was established after ProCyon’s knowledge cutoff date [42] (Supplementary Notes 2, 14). As we refine the prompt with more detailed information—starting with basic phenomena, then including mechanistic details, and lastly adding precise function—the retrieval percentile of STING improves by 6.1 and 16.8 on average, respectively (Figure 2h). To probe which text features drive ProCyon’s retrieval of STING, we ablate key text features from the input prompt (Extended Data Figure 8, Supplementary Figure 6). Removing the function of “autophagy” from the prompt reduces STING’s percentile by 9.7. Replacing both “autophagy” and “ferroptosis” functions with “cell death” recovers STING’s original retrieval.

### Generative text capabilities of ProCyon

We assess ProCyon-SPLIT’s QA capabilities by constructing benchmarks from the test split of ProCyon-INSTRUCT (Figure 3a). Our benchmarks span 115,636 protein-phenotype pairs across 6,153 phenotypes. Comparing to models that can perform protein-phenotype prediction or question-answering (ProtLLM [30]), ProCyon-SPLIT is the best-performing model in 4 out of 6 tasks, and has the the highest performance across tasks with an average accuracy of 74.9% compared to the next-highest accuracy of 63.5% by ESM3-kNN (Figure 3b, Supplementary Table 7, Supplementary Figure 7). ProCyon-SPLIT generalizes to rare and unseen phenotypes, as demonstrated by its QA accuracy across ProCyon-INSTRUCT’s shot-level splits (Figure 3c).

**Figure 3.**
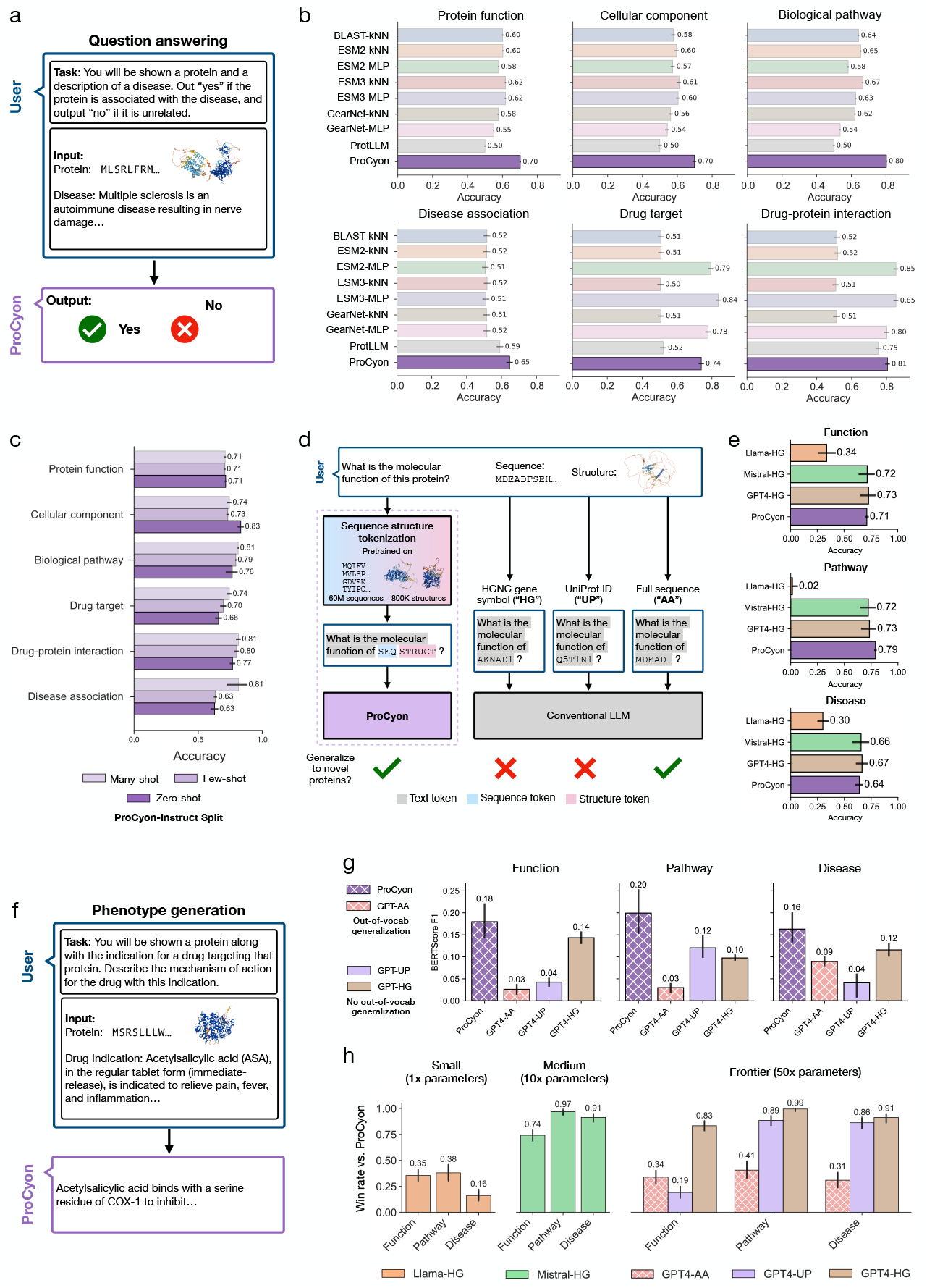
ProCyon generates accurate responses and phenotype descriptions from multimodal prompts. **a)** In QA, ProCyon receives a multimodal prompt (protein + phenotype description) and outputs a yes/no answer. **b)** Bench-marking QA accuracy across models and knowledge domains. Error bars show bootstrapped 95% CIs. **c)** ProCyon’s QA accuracy across different ProCyon-INSTRUCT splits. **d)** ProCyon tokenizes protein sequences and 3D structures using a multimodal encoder; LLMs must use text surrogates, many of which are not generalizable (gene symbol, UniProt ID). **e)** QA performance comparison between ProCyon and LLMs using HGNC gene symbols. **f)** In phenotype generation, ProCyon processes multimodal inputs (proteins, natural language, small molecules) to output open-ended phenotype descriptions. **g)** Phenotype generation benchmark using reference text similarity (mean BERTScore *F*_1_). Hatching indicates ability to generalize to unseen proteins. **h)** Phenotype generation benchmark using LLM-based judging. Win rate refers to preference over ProCyon; models grouped by parameter count.

We compare ProCyon-SPLIT’s QA and phenotype generation capabilities against LLMs and benchmark three text-based representations of proteins: HGNC gene symbol (“-HG”), UniProt ID (“-UP”), and the protein’s full amino acid sequence (“-AA”) (Figure 3d, Methods Sec. 4.6). For QA, we provide the LLMs with the same prompt as ProCyon but replace sequence and structure tokens with the HGNC gene symbol. ProCyon-SPLIT performs comparably to significantly-larger models, such as Mistral Large (Mistral-HG) and GPT-4 (GPT4-HG), and far outperforms similarly-sized Llama3-8B model (Llama-HG) (Figure 3e).

For phenotype generation, where ProCyon-SPLIT generates descriptions of proteins (Figure 3f), we evaluate performance on proteins stratified by UniProt annotation scores (Methods Sec. 4.6), resulting in 657 proteins with reference annotations in three knowledge domains (Function, Pathway, Disease). We first measure quality of the generated phenotypes using BERTScore F1 score [43] against ground-truth annotations (Methods Sec. 4.6). Across all knowledge domains, ProCyon outperforms GPT-4o with any choice of text-based representation for proteins (Figure 3g). We further adopt the LLM-as-a-judge evaluation approach [44], using Claude-3.5-Sonnet [45] to score generated phenotype descriptions (Methods Sec. 4.6, Supplementary Note 15). ProCyon-SPLIT’s open-ended text generation surpasses similarly-sized LLMs (i.e., 1x model parameters) but underperforms significantly-larger models (i.e., 10x and 50x model parameters) (Figure 3h). However, these LLMs are sensitive to the choice of text-based representation, and ProCyon’s outputs are consistently preferred over LLMs when the amino acid sequence (“-AA”) is used.

### Identifying drug-binding protein domains

We investigate ProCyon’s retrieval of domains within proteins that are bound by drugs [46]. Miglitol, a small molecule drug, binds to Maltase-glucoamylase (MGAM), a digestive enzyme with 2,753 amino acids. We use ProCyon to retrieve domains in MGAM by providing ProCyon with a multimodal prompt of miglitol and its mechanism of action. We then score retrieval results over the nine domains in MGAM (Figure 4a, Methods Sec. 4.7). ProCyon correctly identifies the binding domain as the top-predicted domain. MGAM contains two main catalytic domains at the N-terminus (NtMGAM) and the C-terminus (CtMGAM). Although miglitol binds to both domains, its inhibitory effects are stronger for NtMGAM [47]. ProCyon correctly ranks the NtMGAM catalytic domain (`PF01055_349`) as the top binding domain, assigning it a 0.027 higher score than the CtMGAM catalytic domain (`PF01055_2111`). This result is notable given the subtle but important differences in catalytic efficiency between these homologous domains [48].

**Figure 4.**
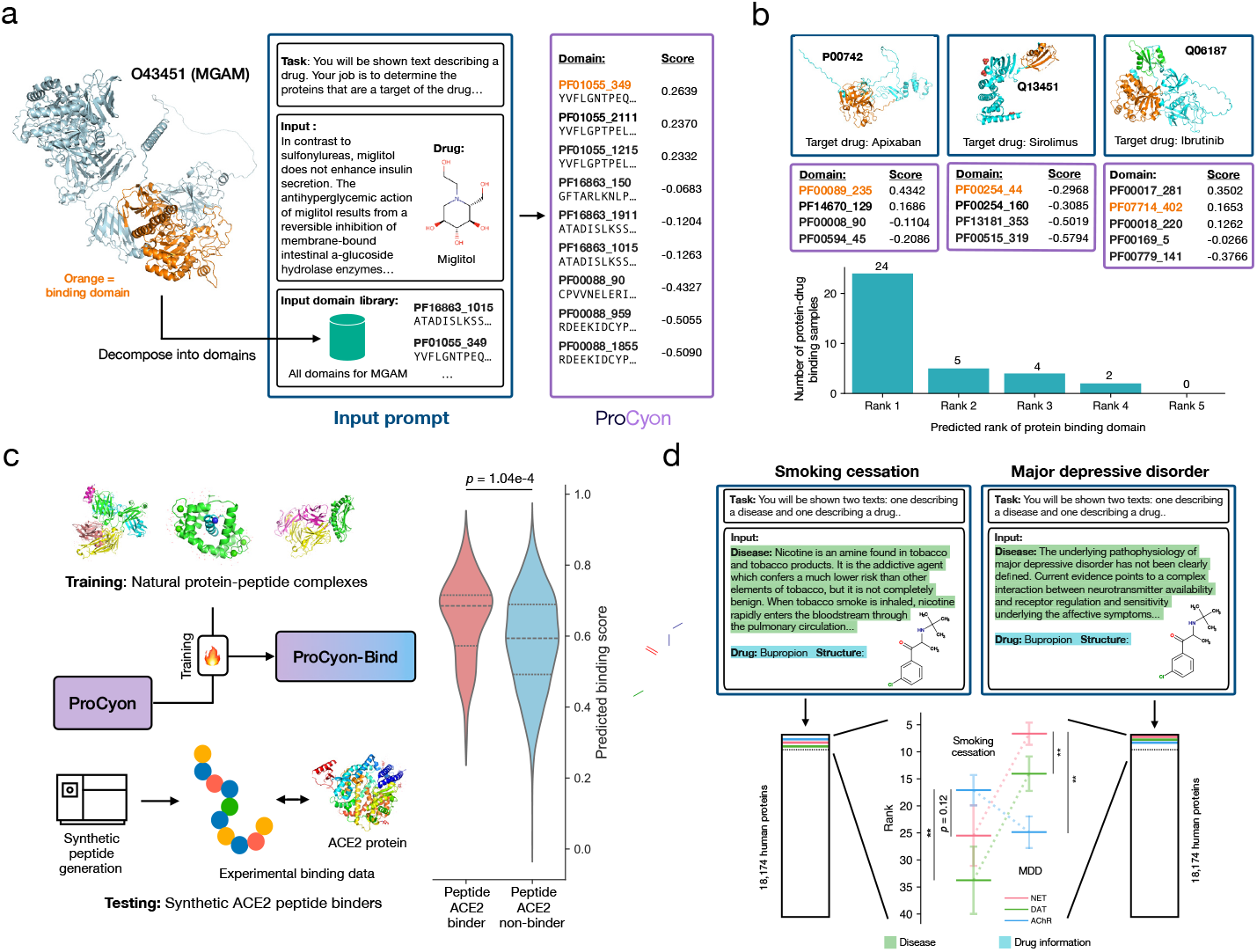
ProCyon models domains, peptides, and small molecules beyond proteins. **a)** ProCyon predicts the small molecule binding domain on proteins, where the input is the description and molecular structure of a drug, and the output is a ranked list of domains on a target protein. In this example, ProCyon retrieves domains of MGAM given the drug miglitol and correctly prioritizes the binding domain for miglitol as the first among the MGAM’s nine domains. Each domain is identified by the Pfam ID and the index of the first amino acid in the domain. **b)** ProCyon ranks the correct drug-binding domain (highlighted in orange) on respective target proteins highly. In Q06187, the green domain (PF00017 281) denotes the ProCyon-predicted binding domain, but the orange domain (PF07714 402) is the correct binding domain. ProCyon predicts the binding domain as top-ranked in 24 examples, second-ranked in 5 examples, and so on. **c)** ProCyon can be finetuned to predict protein-peptide binding. Left, ProCyon is first finetuned on naturally occurring protein-peptide complexes from the Protein Data Bank (PDB), then tested on an experimental dataset of synthetic peptides screened for binding to the ACE2 protein. Right, the predicted binding scores for binders versus non-binders. Two-sided Mann-Whitney U test. **d)** Leveraging ProCyon to perform indication-specific drug target retrieval. In this example, the input consists of a novel task definition, the description of indications (nicotine addiction or major depressive disorder), and the name and structure of bupropion. ProCyon ranks known targets (norepinephrine transporter (NET), dopamine transporter (DAT), and nicotinic acetylcholinergic receptor (AChR)) of bupropion among the top 40/18,174 human proteins. Shown are 95% confidence intervals derived by perturbing words in the input prompt. ** *p*-value *<* 0.001; two-sided Wilcoxon signed rank test.

We extend these analyses to additional protein-drug complexes [46] (Figure 4b). In 68.6% examples of multi-domain proteins, ProCyon ranks the correct drug-binding protein domain at the top of the list. ProCyon outperforms ProtST in this direct setting (Supplementary Figure 8).

### Identifying protein-binding peptides

We evaluate ProCyon’s performance in predicting protein-peptide binding (Figure 4c). We fine-tune ProCyon on physical protein-peptide binding data from PDBBind [49], yielding ProCyon-BIND. We test ProCyon-BIND on an experimentally-measured peptide binding dataset for angio-tensin-converting enzyme 2 (ACE2) [50], a key cardiovascular disease biomarker and SARS-CoV-2 target. This dataset includes 58 detected ACE2 peptide binders and 5,072 non-binders [51]; the imbalance creates a challenging retrieval setting that mimics lead discovery in practice. Despite a small fine-tuning dataset (approx. 1,000 samples), ProCyon-BIND achieves a statistically significant separation between binders and non-binders (*p* = 1.04 *×* 10^*−*4^; Mann-Whitney U test), yielding an AUROC of 0.648 on peptide binders (Figure 4c), compared to the 0.393 AUROC achieved by ESM2 (Supplementary Note 16).

### Indication-specific retrieval of drug targets

ProCyon can also handle queries that cross knowledge domains via its natural language interface. Bupropion is a small molecule drug indicated for major depressive disorder (MDD) and nicotine addiction [52, 53]. Bupropion’s antidepressant effect is attributed mainly to inhibition of the norepinephrine and dopamine transporters (NET, P23975; DAT, Q01959) [54,55], while its putative MoA targets in smoking cessation include the nicotinic acetylcholinergic receptor (AChR, P32297) alongside NET and DAT [56, 57].

We design a novel inference task where ProCyon is prompted with multimodal prompts consisting of bupropion structure and a description of either MDD or smoking cessation (StatPearls [58, 59]; Supplementary Note 17) and asked to retrieve bupropion targets involved in the treatment of either indication. For the smoking cessation prompt, ProCyon ranks AChR at 17.1 on average, significantly higher than NET (25.5) and DAT (33.8) (Figure 4d). Conversely, for the MDD prompt, ProCyon ranks NET (6.7) and DAT (14.1) significantly higher than AChR (24.9), which is aligned with NET and DAT’s involvement in treating MDD. All three proteins (NET, AChR, and DAT) are ranked in the top 40 out of all 18,174 proteins considered.

### Elucidating functions of poorly characterized proteins

We first use ProCyon to functionally annotate AKNAD1 (UniProt: Q5T1N1) (Figure 5a). AKNAD1 is a poorly characterized protein, with no current functional annotation in UniProt and a UniProt annotation score of 3 (out of 5, 5 = best-annotated proteins). AKNAD1 also has low sequence similarity with other human proteins in SwissProt (at most 27% sequence identity (with AKNA) via BLASTp), making functional inference via sequence similarity challenging. ProCyon generates three phenotype descriptions for AKNAD1. Two of these phenotypes are independently validated by experimental evidence in the Human Protein Atlas (HPA) [8] (Methods Sec. 4.10). We also use ProCyon to generate phenotype descriptions for LYRM2, ANKDD1A, and UQCC4 (Figure 5b). ProCyon’s generated phenotype descriptions of LYRM2 in mitochondrial complex I and UQCC4 in the cytochrome b-c1 complex are consistent with UniProt annotations that were added after ProCyon’s knowledge cutoff date. ProCyon’s generation that ANKDD1A inhibits cell proliferation is supported by evidence that its restored expression suppresses glioblastoma growth and invasion [60].

**Figure 5.**
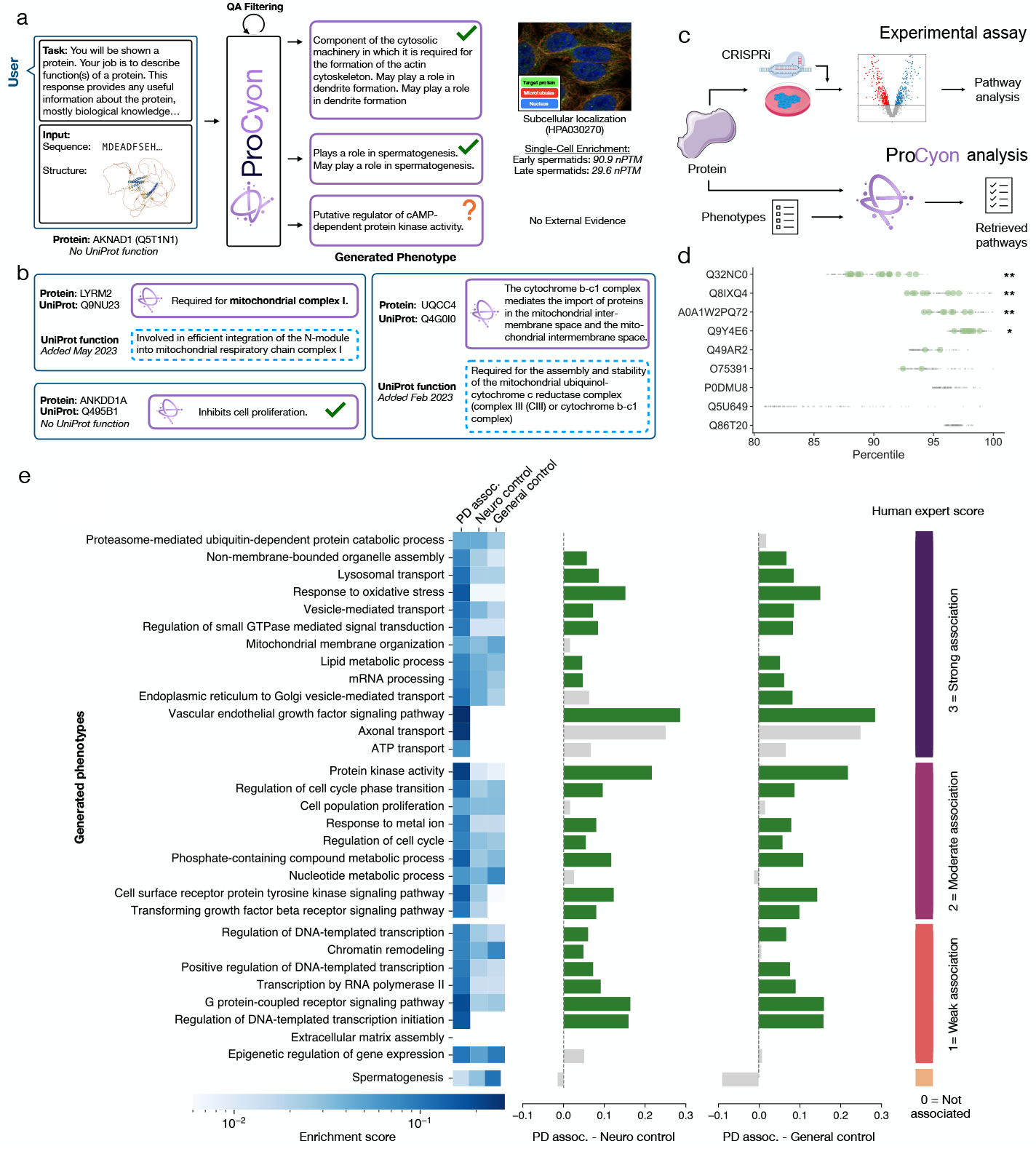
ProCyon characterizes poorly-annotated proteins and their functions. **a)** ProCyon generates phenotype predictions for AKNAD1. QA filtering selects the highest-confidence outputs; two of the three predicted phenotypes are experimentally supported. **b)** Additional ProCyon-generated phenotypes for other poorly characterized proteins. All proteins either have no UniProt-annotated function or have no UniProt function as of the ProCyon knowledge cutoff date. **c)** External evaluation of ProCyon using perturbation datasets. Parallel pathway analysis on genetic perturbation data is used to validate ProCyon-predicted functions for uncharacterized proteins. **d)** Top 100 functions prioritized by ProCyon for each protein. Percentile refers to the percentile of that protein among all 18,174 human proteins in our database for the specific function query. Green dots indicate overlap with perturbation-derived pathways; grey dots indicate other predictions. ** *p <* 0.005, * *p <* 0.05; hypergeometric test. **e)** ProCyon-generated pathways for poorly characterized proteins associated with Parkinson’s disease (PD). Left, heatmap of enrichment scores for each pathway compared to established PD-related proteins (“PD assoc.”), proteins expressed in the nervous system but not in PD or other neurodegenerative conditions (“Neuro control”), and proteins not expressed in brain tissues (“General control”), respectively. Right, differences in enrichment scores between “PD assoc.” and “Neuro control” and “General control”. Statistically significant differences shown in green. Pathways are grouped by PD association score provided by human experts (right).

We use ProCyon’s retrieval capability to predict the functions of nine poorly characterized proteins (Methods Sec. 4.11, Supplementary Figure 9). To validate the predicted functions, we use genome-wide perturb-seq data [61] and perform gene set over-representation analysis (Enrichr [62]) on differentially expressed genes (DEGs) upon gene knockdowns (Figure 5c, Methods Sec. 4.11). ProCyon’s top-100 predicted functions for the Q32NC0, Q8IXQ4, A0A1W2PQ72, and Q9Y4E6 proteins are significantly enriched in the perturb-seq dataset (Figure 5d, Supplementary Figure 10). C18orf21’s (Q32NC0) involvement in rRNA processing and ribosome biogenesis is biochemically validated [63, 64], which is aligned with ProCyon’s predictions (e.g., “rRNA processing”, “rRNA metabolic process”, “ribonucleoprotein complex biogenesis”). MSANTD7 (A0A1W2PQ72) is physically enriched at DNA damage lesions [65], partially aligned with ProCyon’s predictions (e.g., “regulation of DNA binding”, “regulation of intrinsic apoptotic signaling pathway”, “DNA damage response, signal transduction by p53 class mediator”). Further, we examine ProCyon’s capability to annotate missense variants in proteins (Supplementary Note 18, Supplementary Figure 11). We find that by examining changes in generated captions from pathogenic-to-benign variants, ProCyon shows promise to annotate functional effects of protein variants.

We further evaluate ProCyon’s predicted functions for poorly characterized proteins linked to Parkinson’s disease (PD). We select 35 proteins from OpenTargets [66] with reported associations to PD but no presence in ProCyon’s training data, and refer to these proteins as the “PD uncharacterized” set. As a positive control, we compile 962 proteins with validated PD associations (“PD assoc.”) from OpenTargets. We also collect two negative controls: (1) proteins expressed in the nervous system but not associated with PD or neurodegeneration (“Neuro control”), and (2) proteins not expressed in brain tissue (“General control”) (Supplementary Note 19). For each protein set, we use ProCyon to generate phenotype descriptions and then group descriptions into functions using a combination of semantic similarity-based clustering and manual curation. We calculate the overlap in generated functions between PD uncharacterized and PD associated sets, as well as Neuro and General negative controls, using a one-sided proportion z-test (Methods Sec. 4.12). A neurologist/PD specialist scores predicted functions for proteins in the PD uncharacterized on a scale of 0-3 (0=no PD association and 3=strong association, Figure 5e, Extended Data Figure 9). Predicted functions for PD uncharacterized proteins are significantly enriched in the PD associated proteins relative to both negative controls, and are generally given higher scores by the neurologist/PD specialist. Further, most of ProCyon’s predicted functions with neurologist-assigned scores higher than 0 are significantly enriched compared to Neuro and General controls (Figure 5e, Supplementary Note 20).

### Experimental validation in multiple sclerosis brain samples

We integrate ProCyon within a prospective, pathology-guided workflow to nominate multiple sclerosis (MS) genes from post-mortem cortex and probe mechanisms. First, neuropathologists prompted ProCyon using five MS neuropathological hallmarks (Supplementary Table 8). Next, we dissect the superior frontal gyrus (SFG) from the brains of 225 clinically and neuropathologically confirmed MS donors from the Imperial College Tissue Bank, perform neuro-pathological assessment, and conduct bulk RNA sequencing of the brain samples (Figure 6a-f). After data processing, we have 196 samples from which we derive DEGs related to the five MS hallmarks (Supplementary Note 21, Supplementary Figure 12). ProCyon’s average retrieval percentiles of the significant DEGs (FDR corrected *p*-value *<* 0.05) (Figure 6c) are 95.01 *±* 2.34, 85.88 *±* 4.40, 87.15 *±* 8.70, 76.16 *±* 3.46 and 88.18 *±* 5.93 for HLA+ in grey matter, SFG white matter lesions, presence of active lesions, meningeal and perivascular infiltration and HLA+ in white matter, respectively.

**Figure 6.**
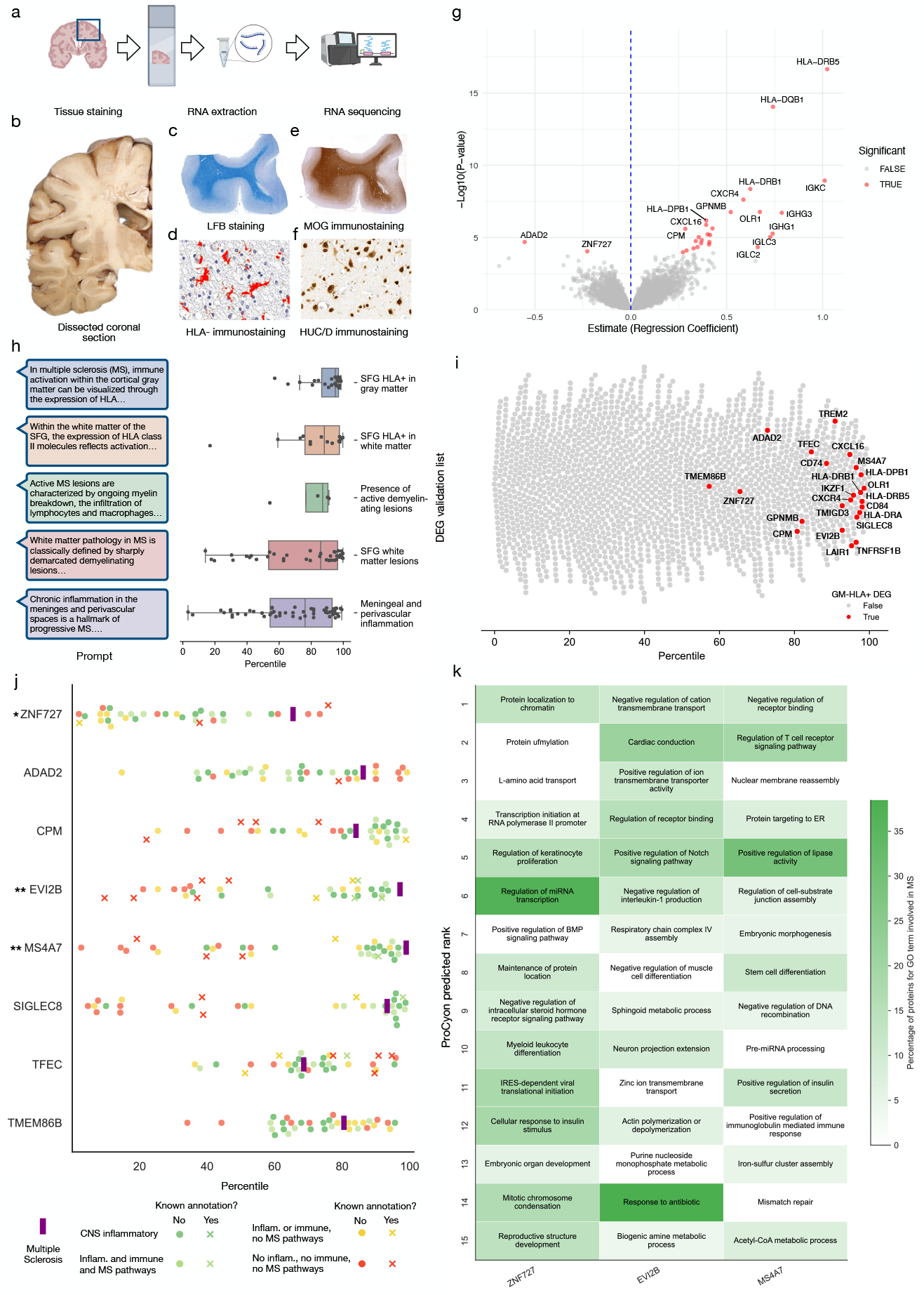
Experimental validation in multiple sclerosis using post-mortem brain RNA-seq with mechanistic characterization by ProCyon. **a)** MS brain tissue was dissected, stained, and RNA-sequenced. **b)** Blocks from the superior frontal gyrus (SFG) were sampled. **c)** Luxol fast blue/cresyl violet distinguished grey and white matter. **d)** MOG staining identified activated microglia/macrophages. **e)** HLA-D^+^ areas were masked in red. **f)** Neurons were labeled with HuC/D. **g)** Differential expression analysis identified HLA^+^-associated genes (FDR *<* 0.05, shown in red). **h)** Validation of ProCyon rankings across five MS hallmarks. Left: retrieval prompts. Right: DEG percentiles for each hallmark. Boxplots show interquartile range and median. **i)** ProCyon-assigned ranks for HLA^+^ DEGs (red) compared to 2000 random non-DEGs. **j)** ProCyon retrieval percentiles for ‘unexpected’ proteins (Methods Sec. 4.13) across control diseases versus MS. Control diseases vary in similarity to MS (bottom legend). X markers denote known protein–disease associations. MS is marked by a vertical purple bar. * = significant compared to MS control group; ** = significant across three control groups (Methods Sec. 4.13). **k)** Top 15 ProCyon-predicted pathways for ZNF727, EVI2B, and MS4A7. Highlighted by the percentage of MS-associated proteins (per GO) in each pathway.

We assess whether ProCyon can identify previously unreported MS genes in human post-mortem cortex. For that, we take HLA-associated DEGs and filter out those previously linked to MS or annotated with inflammatory- or immune-related Gene Ontology terms, yielding eight candidate MS genes: ZNF727, ADAD2, CPM, EVI2B, MS4A7, SIGLEC8, TFEC, and TMEM86B. To assess the disease specificity of each candidate, we use ProCyon to retrieve genes based on their association with MS and related disorders, which are grouped into four categories based on similarity to MS (Methods Sec. 4.13, Supplementary Table 9). We conduct a permutation test using three control gene sets: (i) non-DEG, non-inflammatory, non-immune genes, (ii) non-DEG, non-inflammatory, non-immune, non-MS genes, and (iii) non-DEG, non-inflammatory, non-immune, MS genes. For each control, we calculate statistical significance as the fraction of control genes with higher rankings for MS compared to ProCyon’s candidate genes. EVI2B, MS4A7, and ZNF727 rank as the top three and show significant MS enrichment (*p*-value *<* 0.05). EVI2B and MS4A7 are enriched across all three controls. ZNF727 is enriched only relative to the MS-gene control (Supplementary Table 10).

To probe the mechanism, we prompt ProCyon to identify functions for these three genes and explain each gene’s association with MS pathophysiology (Methods Sec. 4.13, Supplementary Figure 9). Human expert neurologist assessment confirmed that ZNF727 governs chromatin structure, RNA polymerase II activity, and miRNA transcription, all of which are key regulatory processes impaired in inflamed cortical neurons and oligodendrocytes; EVI2B is involved in TCR signaling and cytokine regulation, consistent with its up-regulation in lesion-associated microglia; and MS4A7 both immune functions and lipid metabolism, suggesting its CNS expression and role in sustaining inflammatory and demyelinating processes (Supplementary Note 22).

## Discussion

We present ProCyon, a multimodal foundation model that characterizes protein phenotypes across knowledge domains and scales by using natural language as a unifying modality. PROCYON outperforms approaches for protein-phenotype prediction and generalizes to proteins, phenotypes, and modeling tasks that the model did not encounter during training. We show that ProCyon handles complex, multimodal prompts, enabling tasks such as pleiotropic protein retrieval, indication-specific retrieval of drug targets, and protein-peptide binding prediction. Furthermore, ProCyon generates open-ended phenotype descriptions for poorly characterized proteins, which we validate via external experimental datasets, human expert evaluation, and analysis of previously unpublished transcriptomic data of post-mortem human brain samples.

There are several limitations of ProCyon, which also open up opportunities for future improvement. ProCyon’s conversational capabilities currently lag behind what users have come to expect from interactions with LLMs. Future ProCyon models could incorporate larger-scale LLMs and additional training data to enhance the consistency and coherence of text generation and enable multi-turn interactions for iterative hypothesis refinement. Future datasets could also expand protein representation, for example, by incorporating proteins and annotations from other species or curating additional data sources that provide finer-grained information such as tissue-specific or cell type-specific annotations. A related limitation is ProCyon’s static training dataset, which is unable to leverage continually generated biological research; a potential solution is integrating new information via continual learning, retrieval-augmented generation, or integration with agentic workflows. ProCyon can also be integrated with other data modalities (e.g., transcriptomics, epigenomics) to model protein phenotypes in new biological contexts. Additional modality-specific encoders can be incorporated into ProCyon’s modular design to form a general biological reasoning system.

We envision that models like ProCyon could unify biological modalities across scales to facilitate hypothesis generation, experimental design, and exploration of phenotypic complexity across biological systems. Beyond function querying, ProCyon’s protein-phenotype representations could be adapted for generative protein design (e.g., guiding diffusion models to produce novel functional proteins). ProCyon demonstrates the potential of bridging molecular data and text-based phenotype descriptions to study protein function and human biology.

## Acknowledgements

We thank Jessika Baral, Shanghua Gao, Thomas Wood, and Manuel A. Friese for their helpful discussions on the manuscript and their input on the applications of the model. We are grateful to Max Shad and Ben Sabath from the Research Engineering team at the Kempner Institute for the Study of Natural and Artificial Intelligence at Harvard University for their guidance on multi-node GPU training and large-scale computational analyses. We thank Nathan Palmer and Francis Yu from Harvard Medical School AI Development Central. We also thank Kirill E. Medvedev, Dustin Schaeffer, and Nick V. Grishin for their help in processing data from Drug-Domain on drugs and small molecules bound to domains. We gratefully acknowledge the support of NIH R01-HD108794, NSF CAREER 2339524, US DoD FA8702-15-D-0001, ARPA-H BDF program, awards from Chan Zuckerberg Initiative, Bill & Melinda Gates Foundation INV-079038, Aligning Science Across Parkinson’s (ASAP) Initiative, Amazon Faculty Research, Google Research Scholar Program, AstraZeneca Research, Roche Alliance with Distinguished Scientists, Sanofi iDEA-iTECH, Pfizer Research, John and Virginia Kaneb Fellowship at Harvard Medical School, Biswas Computational Biology Initiative in partnership with the Milken Institute, Harvard Medical School Dean’s Innovation Fund for the Use of Artificial Intelligence, Harvard Data Science Initiative, and Kempner Institute for the Study of Natural and Artificial Intelligence at Harvard University. O.Q. was supported by a graduate fellowship award from Knight-Hennessy Scholars at Stanford University. V.G. was supported by the UK Medical Research Council, MR/W00710X/1. K.H. was supported by Joachim Herz Stiftung via Add-on Fellowships for Interdisciplinary Life Science and the Helmholtz Association under the joint research school “Munich School for Data Science.” Any opinions, findings, conclusions or recommendations expressed in this material are those of the authors and do not necessarily reflect the views of the funders.

## Data availability

Project website for ProCyon is at https://zitniklab.hms.harvard.edu/PROCYON. All processed data supporting the findings of this study are available at https://huggingface.co/datasets/mims-harvard/PROCYON-Instruct. Source data for Figures 1-5 are available in the manuscript. For analyses in Figure 6, fully informed consent for all the tissue and clinical information was obtained via a prospective donor scheme with approval from the Multicentre Research Ethics Committee (MREC/02/2/39).

## Code availability

We release all of ProCyon’s model weights and codebase (https://github.com/mims-harvard/PROCYON) as well as an online demonstration of ProCyon such that the research community can build on and explore our model’s capabilities.

## Authors contributions

O.Q., Y.H., M.K., and M.Z. conceived the study. O.Q., Y.H., R.C., V.G., Y.E., M.K., and M.Z. designed the experiments. O.Q., Y.H., R.C., V.G., G.D., L.T., T.Co., and K.H. collected and processed the data. G.A, O.H., I.F., D.O., and R.N. collected the multiple sclerosis experimental data. O.Q., Y.H., R.C., T.Ch., G.D., and T.Co. developed the model and training codebase. O.Q., Y.H., R.C., V.G., L.T., Y.E., G.A., and A.N. conducted the experimental analyses and interpreted the results. T.Ch., G.D., A.N., J.B., T.H., B.P., J.Z., R.N., F.J.T, and V.K. provided feedback on the analyses. O.Q., Y.H., R.C., V.G., M.L. and M.Z. wrote the manuscript with input from all co-authors. M.K. and M.Z. supervised the research, with M.Z. leading the study.

