## Extended Data Figures and Methods for "ProCyon: A multimodal foundation model for protein phenotypes"

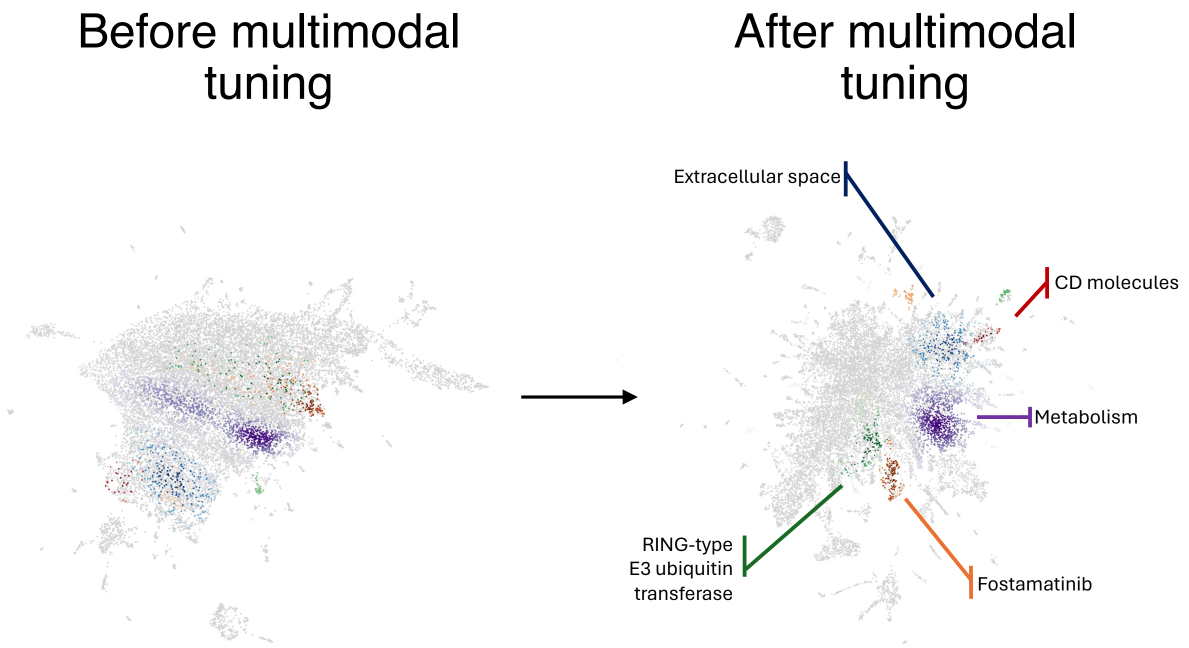


**Extended Data Figure 1: Multimodal retrieval space with ProCyon.** Density of proteins with the same annotation across multiple databases before and after multimodal tuning. The annotations with the highest or second highest number of proteins per database are visualized in the figure. The databases included are: Reactome (purple), EC (green), GTOP (red), GO (blue) and DrugBank (orange). The annotations visualized are: Fostamatinib targets (Drugbank), Metabolism (Reactome), CD molecules (GTOP), Extracellular space (GO), and RING-type E3 ubiquitin transferase (EC). The left embedding was generated using ESM-2 embeddings and corresponds to a sequence-only embedding space (also the initial embedding space of ProCyon). The right embedding figure was generated using the ProCyon embeddings and corresponds to the multimodal tuned embedding space after the ProCyon training process.

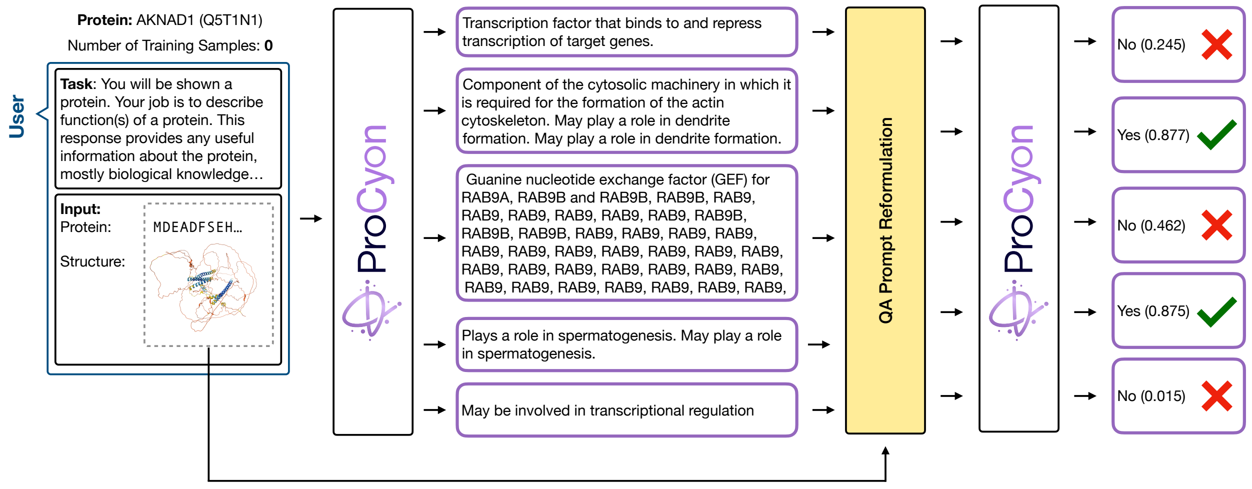


**Extended Data Figure 2: QA filtering with ProCyon.** QA filtering for the example shown in Figure 5a. ProCyon originally produces $N$ texts, and then we reformulate the output description as a QA query and pass it back to the model, getting output yes/no scores (as described in Section 2.3). Note that this procedure removes incoherent text (such as middle example with "RAB9" repeats) and vague general text ("May be involved in transcriptional regulation") when necessary. Since QA filtering also leverages an internal and explicitly-trained mechanism of ProCyon, it also aids in removing biologically-infeasible texts.

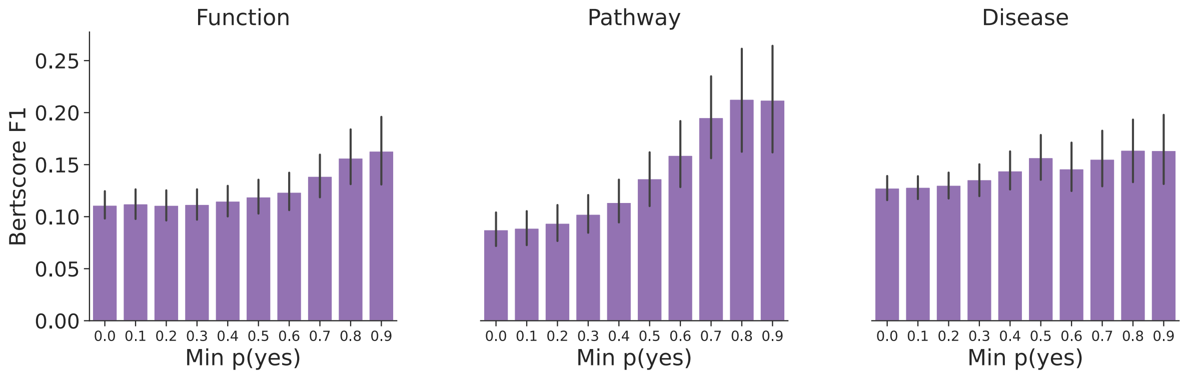


**Extended Data Figure 3: QA filtering improves caption quality.** Shown are BertScore F1 scores with higher QA filter thresholds ("Min p(yes)"). As the threshold increases, the quality of captions improve. This holds across multiple knowledge domains, and QA filtering seems to be more effective for function and pathway domains.

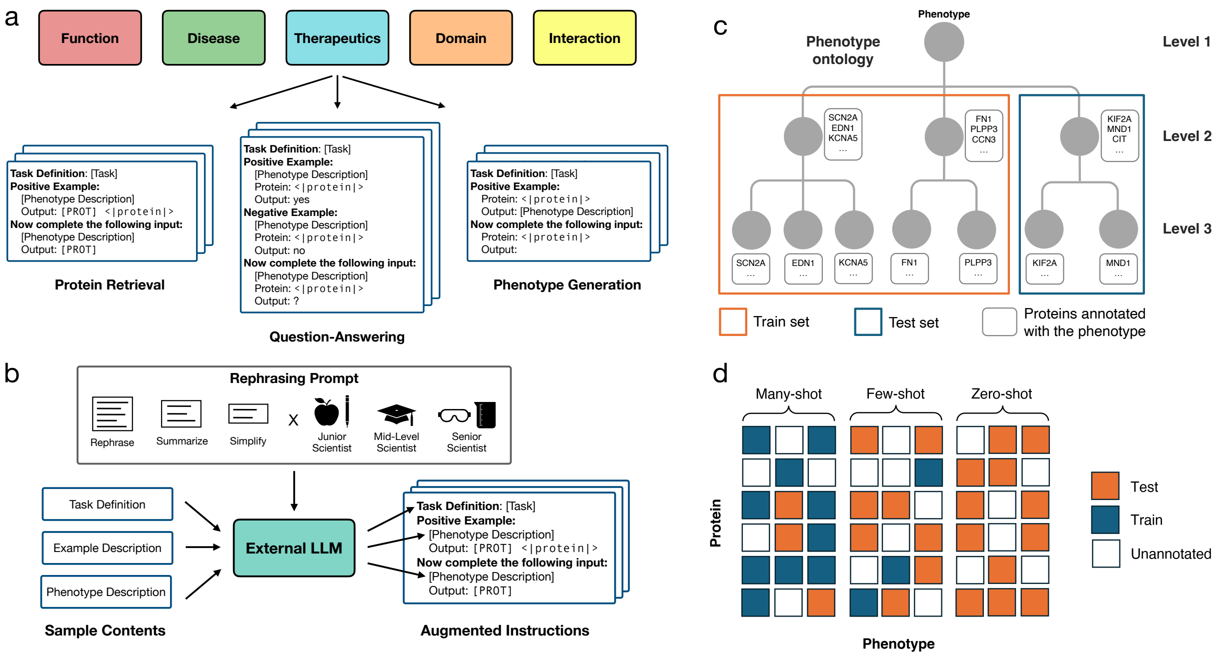


**Extended Data Figure 4: Building ProCyon-Instruct.** **a)** Instruction tuning templates for protein-text pairs from each database in ProCyon-Instruct. Samples in each database can be transformed into an instruction input via the presented instruction templates that formulate protein-text pairs into either a retrieval prompt, a question-answering prompt, or a phenotype generation prompt. **b)** Pipeline for rephrasing descriptions and definitions. Using the templates shown in (a), we take the Task Definition, Example Description ("phenotype description" after the positive and negative examples), and Phenotype Description, input them to an external, proprietary LLM (GPT-4o), and then insert these rephrased versions into the instructions. Note that only the retrieval instruction is shown, but the same process is repeated for all three types of tasks. We prompt the LLM with a rephrasing level and expertise level as discussed in Methods Sec. 3.1. **c)** Illustration of our dataset splitting strategy for each database. To minimize the direct leakage of information, we utilize the hierarchical ontology of phenotypes (e.g., Gene Ontology, ATC code, Enzyme Commissions numbers), and derive the test set by holding out subtrees of phenotypes, along with all protein associations involving these phenotypes. For diseases (DisGeNet) specifically, we utilize the molecular similarity between diseases (defined as the Jaccard similarity between their respective associated proteins) and hold out all diseases with high molecular similarity to certain "seed" diseases (Methods Sec. 1 for details). **d)** Overview of dataset split across many-, few-, and zero-shot splits.

**Sample Instructions**

**Definition**: You will be shown a protein. Your job is to describe a disease, which could be common, rare (Mendelian), complex or environmental, and this protein should be associated with the disease…

**Positive example 1:**
Context: Transcriptional repressor. Represses transcription from both N box- and E box-containing promoters. May with HES1, cooperatively regulate somite formation in the presomitic mesoderm (PSM). May function as a segmentation clock, which is essential for coordinated somite segmentation (By similarity).
Protein: <|protein|> <|struct|>
Output: [ANSWER] A rare disorder caused by mutations in the DLL3 gene, MESP2 gene, LFNG gene, or HES7 gene. It is characterized by abnormal development of bones in the spine and ribs.

**Now, complete the following instance:**
Context: May be involved in protein trafficking.
Protein: <|protein|> <|struct|> Output: [ANSWER] A major psychotic disorder characterized by abnormalities in the perception or expression of reality. It affects the cognitive and psychomotor functions. Common clinical signs and symptoms include delusions, hallucinations, disorganized thinking, and retreat from reality.

**Knowledge Domain:** Disease
**Dataset:** DisGeNet
**Relation:** All
**Task:** Disease-Gene Association
**Capability**: Phenotype Generation

**Definition**: You will be shown a protein and its functions, along with a drug and its indication. Your job is to describe the mechanism of action of the drug…

**Positive example 1:**
Context: Tyrosine-protein kinase that acts as cell-surface receptor for fibroblast growth factors…
Drug Indication: Erdafitinib is a pan-fibroblast growth factor receptor (FGFR) tyrosine kinase inhibitor that is indicated for the treatment of adult patients with locally advanced or metastatic urothelial carcinoma…
Drug: <|drug|> Protein: <|protein|> <|struct|>
Output: [ANSWER] Erdafitinib is subsequently an oral selective pan-FGFR kinase inhibitor that binds to and inhibits the enzymatic activity of expressed FGFR1, FGFR2, FGFR3, and FGFR4…

**Now, complete the following instance:**
Context: Catalyzes the reversible hydration of carbon dioxide. Can hydrate cyanamide to urea.
Drug Indication: For treatment of chronic open-angle glaucoma and acute angle-closure glaucoma
Drug: <|drug|> Protein: <|protein|> <|struct|>
Output: [ANSWER] Methazolamide is a potent inhibitor of carbonic anhydrase. Inhibition of carbonic anhydrase in the ciliary processes of the eye decreases aqueous humor secretion, presumably by slowing the formation of bicarbonate ions with subsequent reduction in sodium and fluid transport.

**Knowledge Domain:** Therapeutics
**Dataset:** DrugBank
**Relation:** Target
**Task**: Target-specific mechanism of action generation
**Capability:** Phenotype Generation

You are pursuing a MD, MD/PhD, PhD, or ScD in biomedical sciences. You are learning deep, specialized medical science knowledge; you can use specialized expert words, but not too many.

**Definition**: You will be shown text describing a biological process or pathway. Your job is to determine the proteins that are involved in the process or pathway described. Involvement is determined by life scientists, using evidence from the scientific literature and input from experts in each pathway.

**Positive example 1**:
Description: Matrix metalloproteinases (MMPs), formerly known as matrixins, are zinc and calcium dependent proteases…
Protein: <|protein|> <|struct|>

**Now, complete the following instance**:
Description: RNA polymerase II (Pol II) catalyzes DNA-directed mRNA synthesis in gene transcription…
Protein: [PROT]

**Knowledge Domain:** Function
**Dataset:** Reactome
**Relation:** All
**Task**: Pathway annotation
**Capability:** Retrieval

**Definition**: You will be shown two proteins. Your job is to determine if the two protein interact based on experimental data, such as, affinity chromatography. Experimental methods, including yeast two-hybrid, affinity chromatography, tandem affinity purification coupled to mass spectrometry, among many others, establish direct or indirect physical interactions between protein pairs. You may only output yes or no. If the protein shown is the one described, you should respond yes. Otherwise, you should respond no.

**Positive example 1:**
Protein 1: <|protein|> <|struct|>
Protein 2: <|protein|> <|struct|>
Output: [ANSWER] yes

**Negative example 1:**
Protein 1: <|protein|> <|struct|>
Protein 2: <|protein|> <|struct|>
Output: [ANSWER] no

**Now, complete the following instance:**
Protein 1: <|protein|> <|struct|>
Protein 2: <|protein|> <|struct|>
Output: [ANSWER] no

**Knowledge Domain:** Interaction
**Dataset:** STRING
**Relation:** Experiment
**Task**: Experimental protein interaction prediction
**Capability:** QA

**Extended Data Figure 5: Example PROCYON instructions.** Example instructions for four different tasks in PROCYON-INSTRUCT, including disease-gene association, target-specific mechanism of action (MoA) generation, pathway annotation, and experimental protein interaction prediction. These examples show how the various knowledge domains can be represented as a unified instruction template which interleaves multimodal inputs (i.e., proteins, domains, etc.) into the text within each instruction. For a more detailed example, see Supplementary Figure 6.

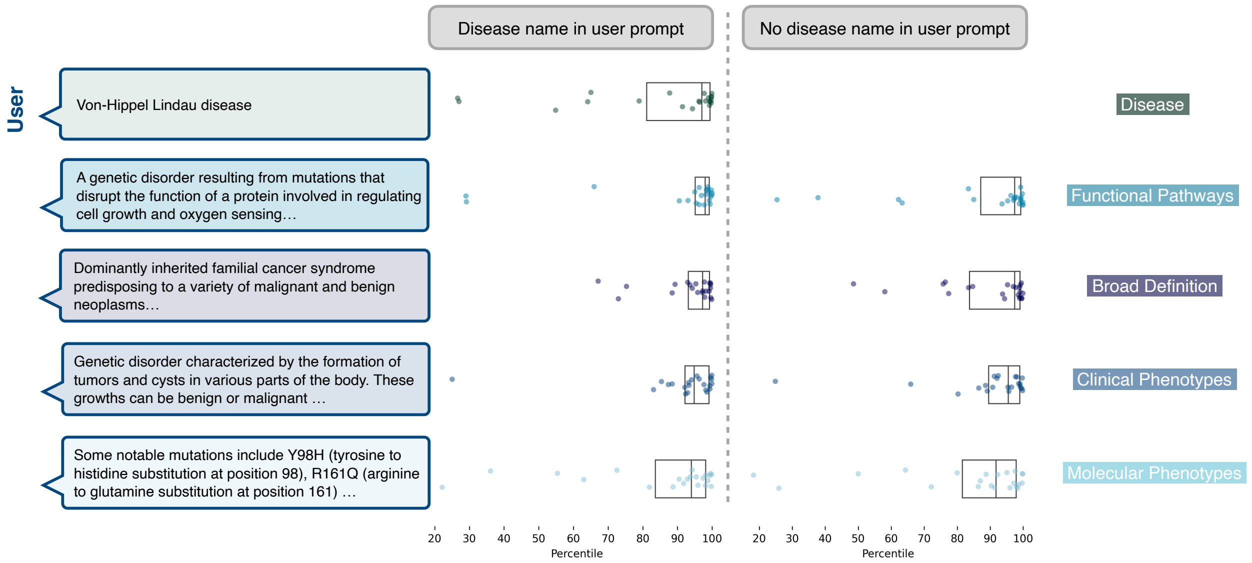


**Extended Data Figure 6: Retrieval on monogenic diseases with PROCYON.** PROCYON can successfully retrieve the causal gene of 22 monogenic diseases using as input prompt descriptions providing different information, and with or without the disease name. The disease descriptions correspond to the disease name, clinical phenotype, functional pathways, molecular phenotype and a broad definition. The causal gene was originally identified experimentally through cloning and linkage analysis.

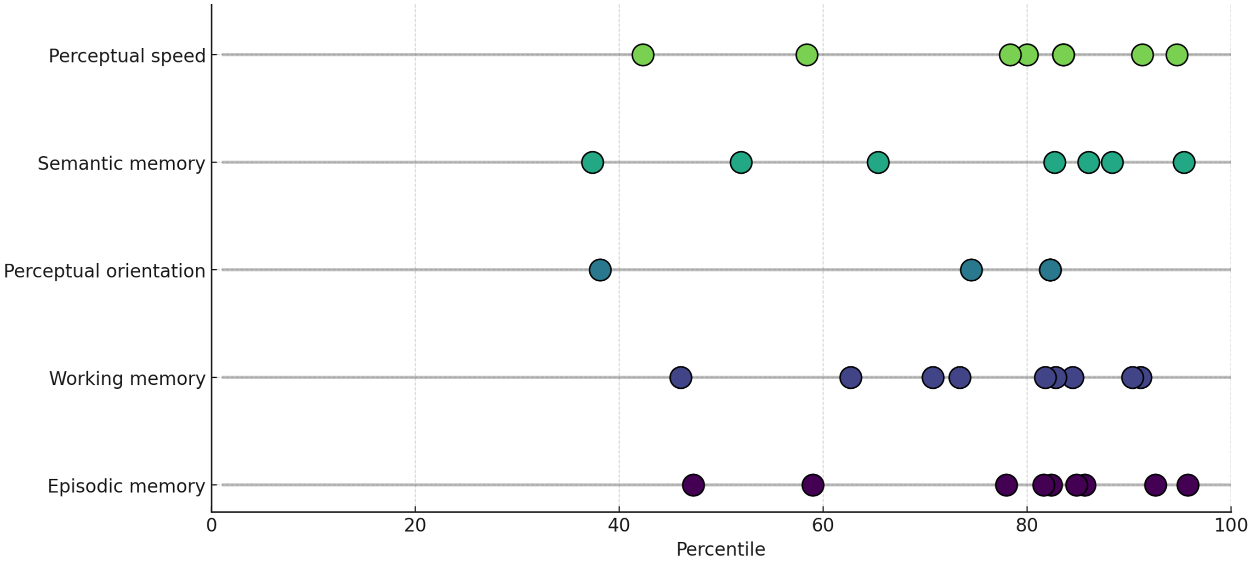


**Extended Data Figure 7: Retrieving genes associated with cognitive domains.** Percentile ranks of genes associated with key cognitive domains commonly affected in dementia, as retrieved by PROCYON. Each line represents a cognitive domain, with highlighted points indicating genes independently identified as differentially expressed in excitatory neuron subtypes [67]. This analysis demonstrates PROCYON 's ability to recover biologically relevant associations absent from its training data and not explained by expression levels alone. Details in Supplementary Note 13.

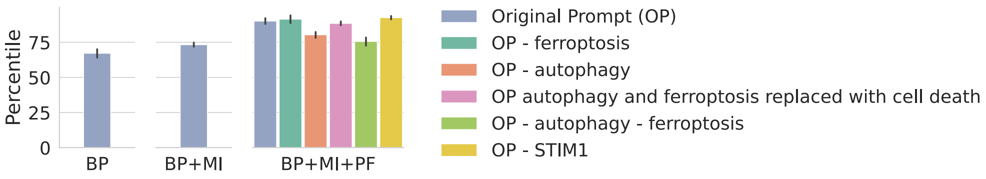


**Extended Data Figure 8: STING prompt ablations.** Percentiles of STING (UniProt: Q86WV6) when querying ProCyon with different levels of evidence about the potential involvement of STING in neuronal inflammatory stress response (BP: Basic Phenomenon; BP+MI: Basic Phenomenon + Mechanistic Insights; BP+MI+PF: Basic Phenomenon + Mechanistic Insights + Precise Function). The evidence descriptions are manually curated based on [42] (Methods Sec. 4.5). The level 3 prompt is ablated to test the sensitivity of ProCyon to the removal of various text features. Full details about ablated prompts are included in Supplementary Note 14.

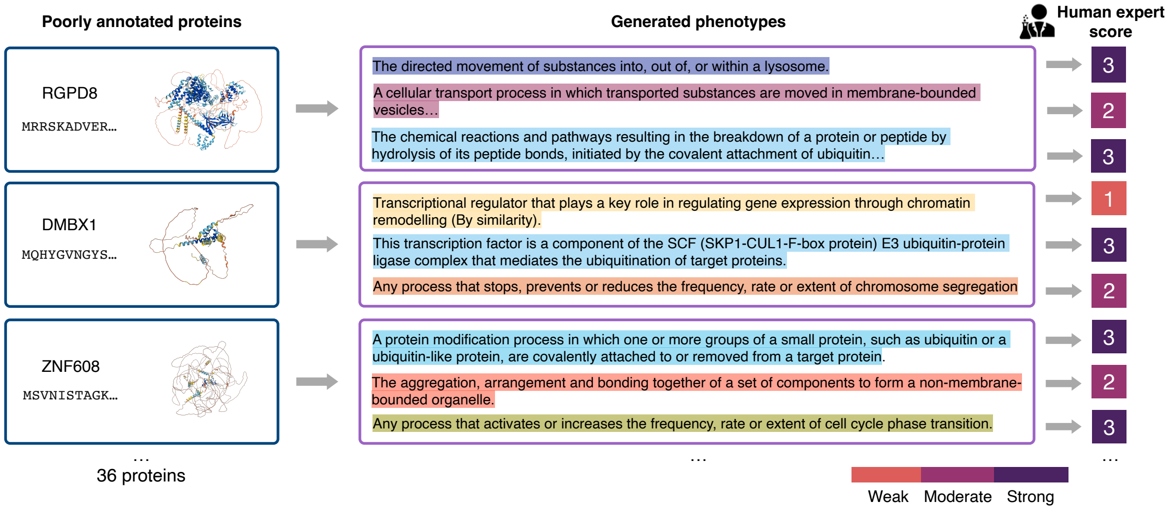


**Extended Data Figure 9:** We designed a study of proteins with known association to Parkinson's disease (PD) but with poor functional annotation in literature. We generated captions with PROCYON, clustered them via manual and automatic methods, manually annotated the pathways identified by each cluster with the help of two independent experts, and provided the pathways to a physician-scientist who specializes in PD and related disorders, who scored their PD association on a scale of 0-3 (with 0 being no association and 3 strong association).

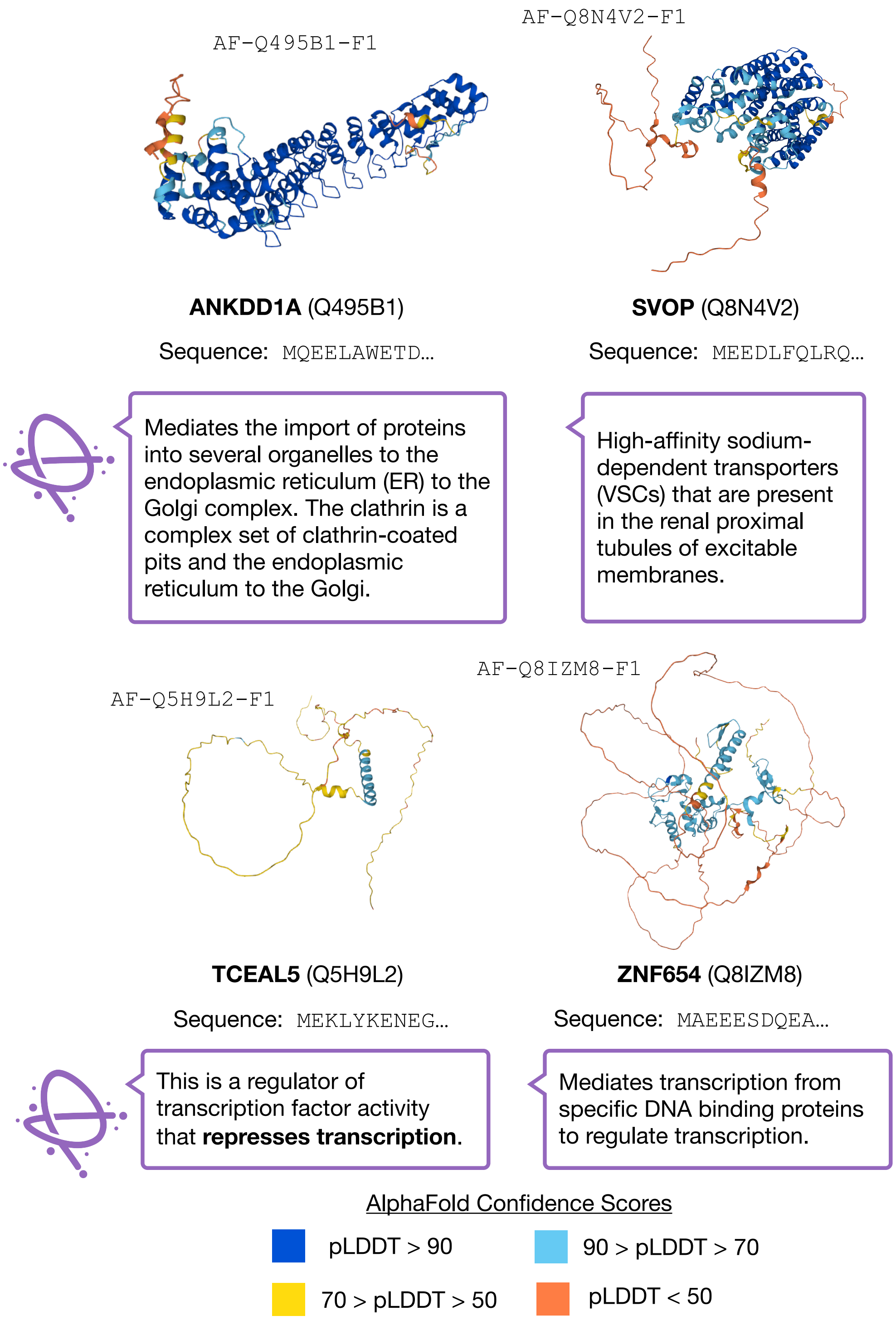


**Extended Data Figure 10:** PROCYON captions for poorly-characterized proteins determined by [68] to be associated to Parkinson's disease via $\alpha$-synuclein toxicity screening. Each protein has no phenotypic relations in PROCYON-INSTRUCT, representing natural zero-shot examples for PROCYON. We show PROCYON descriptions for biological processes the proteins may be involved in. Shown is also the confidence scores for AlphaFold-predicted structures of each protein; this highlights the poor experimental characterization of each protein considered.

### Online Methods

### 1 Datasets

We introduce our dataset ProCyon-Instruct, which contains diverse datasets curated from 12 sources that span biological *knowledge domains*. This dataset is available for download at https://huggingface.co/datasets/mims-harvard/ProCyon-Instruct.

#### 1.1 ProCyon-Instruct dataset

First, to specify the types of datasets in ProCyon-Instruct, we define five different *knowledge domains* into which each dataset can be binned:

**Function.** The function knowledge domain annotates proteins and domains with pathway-level biological information, including molecular functions and biological processes in which each protein is implicated. To curate such annotations, we source data from Gene Ontology [69], Reactome [70], Enzyme Commissions [71], and UniProt [24].

**Protein domains.** This knowledge domain contains data on domain families, which are subunits of proteins that are shared across many proteins and communicate distinct, shared functions. We derive functional information about domains from Gene Ontology [69] and Pfam [72,73].

**Disease.** The disease knowledge domain annotates proteins with diseases they are associated with, where associations are defined based on established evidence linking genetic variations or mutations to the diseases. To curate such annotations, we source data from DisGeNet [74] and Online Mendelian Inheritance in Man (OMIM) [75].

**Therapeutics.** The therapeutics knowledge domain annotates proteins with their associations with drugs, including their involvement in the drug mechanism of action, uptake, and metabolism. To curate such annotations, we source data from DrugBank [76] and IUPHAR/BPS Guide to Pharmacology [77].

**Interaction.** The interaction knowledge domain encompasses interaction between multiple proteins, domains, or peptides. We curate the STRING dataset [78] which includes protein-protein relationships of coexpression, homology, and experimental evidence. Finally, we include a Protein-Peptide binding dataset from PDBBind [49] which encompasses physical interactions between biological macromolecules more broadly.

Detailed information on the downloading and processing of each separate dataset is included in Supplementary Note 2. There are a number of resources we do not include for each knowledge domain. Some exclusions are due to the restricted access to the full repository of data. For some others, exclusions are due to the need of external validation. For example, we utilize Open Targets database for external validation, which is thus not included.

#### 1.2 Dataset splitting for PROCYON-INSTRUCT

All datasets in PROCYON-INSTRUCT, with the exception of protein-protein interaction and protein-peptide binding datasets, are composed of samples of protein- or domain instance-phenotype pairs (in short, "protein-phenotype pairs"). For most datasets, such a pair means an annotation of a textual description of a phenotype to a protein or domain instance. For the DrugBank dataset specifically (protein-drug associations), associations are further stratified into "target", "carrier", "enzyme", and "transporter", representing distinct functional roles of proteins at different stages of a drug's absorption, distribution, metabolism, or excretion within the human body. To robustly evaluate the capability of models to annotate proteins with phenotypes beyond controlled vocabularies, we split protein-phenotype pairs in each dataset into one training set and multiple test sets, evaluating different levels of generalizability. We generally follow the following two steps to split each dataset:

1. First, unique phenotypes from this dataset are split into three different phenotype sets: many-shot, few-shot, and zero-shot, with a ratio of approximately [1] 0.7:0.15:0.15, based on the ontology of phenotypes in the respective source as specified in Supplementary Note 6 to ensure minimal leakage across the train and test sets. Note that there is no notion of "protein" in this step yet, except that we make sure that the few-shot phenotypes are annotated with more than five proteins to ensure sufficient number of samples (protein-phenotype pairs) to be split to train split wherever possible.

2. Then, we split about 80% of all protein-phenotype pairs containing phenotypes in the many-shot phenotype set into the training set, the rest into the many-shot test set. For each phenotype in the few-shot phenotype set, $k$ protein-phenotype pairs are split into the training set ($k$ is set to be either 2 or 5 depending on the average number of associations for phenotypes in each dataset), the rest into the few-shot test set. All protein-phenotype pairs containing

[1] As will be specified in later subsections, it is oftentimes not possible to split the phenotypes with an exact ratio as desired due to the stringency brought by additional requirements in our splitting strategy for those datasets. The exact ratios of splits for each dataset can be found in Supplementary Note 6.

zero-shot phenotypes are kept in the zero-shot test set.

In summary, the many-shot test set contains protein-phenotype pairs with phenotypes that are seen much during training, the few-shot test set contains protein-phenotype pairs where the phenotypes are only seen at most $k$ times during training, and the zero-shot test set contains protein-phenotype pairs with phenotypes that are not seen during training. Three validation sets are also split in a similar manner for model development purpose. In particular, for the few-shot and zero-shot phenotype sets, we split phenotypes further 1:2 (validation:test). Please note that in most descriptions, we omit the mentioning of validation set for simplicity. Training sets from all datasets are merged as the final training set. Finally, we perform additional adjustments to dataset splits via cross-database matches (where available) and semantic similarity of phenotypes (Supplementary Note 7). Details on each specific split are included in Supplementary Note 6.

#### 1.3 Generating dataset rephrasings

Language from curated biomedical databases often consist of boilerplate terminology that is standardized across terms in the database. For example, the Gene Ontology text for “lipoxin A4 biosynthetic process” reads “The chemical reactions and pathways resulting in the formation of lipoxin A4. Lipoxin A4 is a C20 hydroxy fatty acid having (5S)-, (6R)- and (15S)-hydroxy groups as well as (7E)- (9E)-, (11Z)- and (13E)-double bonds.” while the text for “lipoxin B4 metabolic process” reads “The chemical reactions and pathways involving lipoxin B4. Lipoxin B4 is a C20 hydroxy fatty acid having (5S)-, (14R)- and (15S)-hydroxy groups as well as (6E)- (8Z)-, (10E)- and (12E)-double bonds.” Both sentences have roughly the same grammatical syntax, a structure that is represented across much of the Gene Ontology database. These boilerplate descriptions might be comprehensive and detailed enough for experts in the field, but this can make such terminology inconcise and impractical for a broader scientific audience. In addition, this lack of diversity in text is detrimental to language model training, where diversity of the dataset is crucial for learning [79–82]. Therefore, we implement a data augmentation strategy whereby we prompt GPT models [23] to rephrase the entities from each database to represent the diversity of user inputs that the model might encounter in deployment. To further increase diversity, we include prompting that encourages GPT to output descriptions in the style of researchers with different levels of expertise.

To generate such rephrasings, we prompt `GPT-3.5-turbo-0125` to generate responses in a JSON format that includes several definitions of expertise levels. We define three expertise levels: 1) junior scientists, which are individuals at a Bachelor's degree level who have basic medical science knowledge but no specialized biological knowledge, 2) mid-level scientists, which are individuals pursuing a graduate or professional degree in the biomedical field and thus are developing deep knowledge of biomedicine, and 3) senior scientists, who have specialized knowledge of biomedical concepts and have more than five years of postdoctoral research experience. In addition, we define two levels of description transformations 1) rephrasing: an "alternate phrasing" of the phenotype and 2) summarization: a shortened and more concise version of the phenotype. GPT is prompted to generate a "rephrasing" and "summarization" for each expertise level. The full description of the expertise levels are included in Supplementary Note 3. We include the full prompt to GPT in Supplementary Note 23. Finally, we include a table with all fields that were rephrased in Supplementary Table 11. We do not publicly release these rephrased entities; please contact the authors of this work to request access to these augmented descriptions.

### 2 ProCyon Model

**Overview.** We introduce ProCyon, a novel multimodal foundation model that is built on top of transformer deep learning architectures. ProCyon is designed to process multiple biomedical modalities in the input—including protein sequence, protein structure, ligand structure, and natural language—while generating text and prioritizing proteins in the output space. The core of ProCyon is built on decoder-only transformer-based language models, specifically the LLaMA family of models [31,83]. ProCyon is inspired by pioneering architectures in multimodal vision-language models (VLMs), including LLaVA [84], BLIP [35,85], FROMAGe [86], and Flamingo [87], which center around LLMs for processing multimodal information. ProCyon can interleave multimodal inputs into textual prompts, enabling the specification of in-context examples and diverse types of input queries, such as prompting the model for protein-protein interactions. In addition, ProCyon is trained to retrieve and prioritize proteins and arbitary amino acid sequences based on user input queries. In this section, we will describe the exact specifications for our model architecture, which is in supplement to the model implementation given in our codebase https://github.com/mims-harvard/ProCyon. We fully release all of ProCyon's model weights and codebase as well as extensive demos and steps for using ProCyon such that the research community can build on and explore our model's capabilities.

#### 2.1 Capabilities and modes of PROCYON

PROCYON is a foundation model that has a variety of multimodal capabilities. We start this section by describing how a text-only LLM can be made to take multimodal inputs via multimodal token composition, and then we proceed to describe how we can leverage different "modes" of the model to accomplish diverse tasks at inference time. In terms of input and output structures, PROCYON has two main modes: retrieval and text generation. For text generation, we define two main sub-modes, including question answering (QA) and phenotype generation. Details for inference of PROCYON are included in Methods Sec. 2.3, and training details are described in full in Methods Sec. 3.

**Multimodal token composition.** PROCYON is built on top of a pre-trained generative large language model (LLM) which is a decoder-only transformer architecture [88]. The input to the LLM module is a set of token embeddings $\{\mathbf{t}_1, ..., \mathbf{t}_n | \mathbf{t}_i \in \mathbb{R}^{d_{\text{input}}}\}$. The LLM is pretrained on a large text-only corpus, i.e., each $\mathbf{t}_i$ represents a piece of a word as defined by the Byte-Pair Encoding (BPE) tokenizer [89]. We adopt the base tokenizer used by LLaMA-2 and LLaMA-3 for PROCYON-SPLIT and PROCYON-FULL, respectively.

An interleaved multimodal input consists of a combination of text, protein sequence, protein structure, and drug ligand structure inputs, unified through token embeddings. For example for a sequence `sequence_1` with corresponding structure `structure_1`, an interleaved input with text might be "Describe the role of `sequence_1` `structure_1` in obesity". To denote these multimodal inputs in the model, we introduce new input tokens to the tokenizer. <|protein|> denotes a protein sequence input, <|struct|> denotes a protein structure input, and <|drug|> denotes a drug ligand input. During tokenization, we keep track of the indices of each multimodal input token and use this later for inserting multimodal embeddings.

Each input modality gets a specific encoder that has been especially designed with inductive biases to match the given modality. To encode the protein sequences, we utilize a large protein language model which has been pretrained on over 60 million proteins [32]. For protein structures, we use an equivariant geometric deep learning model built to process 3-dimensional molecular structures [19]. For ligand drug structures, we use another geometric deep learning model that achieves state-of-the-art performance for molecular classification tasks [90]. More details on exact specifications of the encoders for each modality are included in Methods Sec. 2.2. During a forward pass, we deconstruct a user input into the respective modalities provided to the model, and we pass each input entity into the encoders for their respective modality; this then gives us embeddings for each multimodal input entity.

The next step is to connect the output spaces of the protein and drug encoders to that of the LLM. We use three-layer multi-layer perceptrons (MLPs) to accomplish this task which we refer to as "Connectors". Specifically, we use the "Token Connectors" to project the embeddings of proteins and drugs into the input space of the LLM. Here, we describe only the token connectors (other connectors are defined in subsequent sections). We define three token connectors: 1) $\mathbf{C}_p^t$ which takes as input the output embedding from the protein language model, 2) $\mathbf{C}_s^t$ which takes as input the protein structure embedding, and 3) $\mathbf{C}_d^t$ which takes the drug/ligand structure as input. The input to each connector matches the dimensionality of the output embedding from each multimodal encoder, and the output from the final layer of the MLP matches the dimensionality of the input token embeddings for the LLM. Once each unified embedding is received, we take these embeddings and pass them to the LLM in the index of the multimodal indicator tokens described previously. Thus, each multimodal entity becomes a kind of "soft token" input to the model, which allows it to flexibly encode a wide variety of modalities that can be interleaved as complex instructions.

**Retrieval mode.** The retrieval mode is defined as a query in which the input is a phenotype and the output is a list of prioritized amino acid sequences, which can be either proteins, domains, or peptides, depending on user preferences. Practically, output is returned to the user as an ordered list of proteins $p_i$ with corresponding retrieval scores $s_i$, i.e., $\{(p_1, s_1), ..., (p_n, s_n)\}$ where $s_i < s_{i-1}$. For retrieval demonstrations, we return the lists in descending order, where the first protein returned is the most relevant according to the model. Inputs can be any combination of phenotype given as either text or other amino acid sequences. Some example tasks that can be formulated as retrieval tasks are disease-gene prioritization, drug target identification, and homologous protein retrieval. See Figure 1a for more examples.

To enable the pre-trained LLM to accomplish this task, we introduce a special token [PROT] that denotes a retrieval task. This is similar to the approach taken by FROMAGe [86], which grounds language model outputs in libraries of images. We then access the hidden representation of the last layer of the LLM that corresponds to the next-token prediction of the [PROT] token, $\mathbf{z}_p^l \in \mathbb{R}^{d_{\text{LM}}}$, where $d_{\text{LM}}$ is the hidden dimensionality of the last layer in the language model. We then define two MLP connectors to aid in the retrieval procedure. First, $\mathbf{C}_p^q$ is known as the Query Connector, and it projects the output embedding from the language model $\mathbf{z}_p^l$ (the query) to a shared retrieval latent space space $\mathbf{z}_r^l \in \mathcal{Z}^r$. Second, we define a Retrieval Connector $\mathbf{C}_p^r$, which projects a set of candidate protein sequences $\mathcal{P}$ into the retrieval embedding space $\mathcal{Z}^r$. This shared latent space is then optimized via contrastive learning (Methods Sec. 3.4) and leveraged for multimodal retrieval (inference procedure described in Methods Sec. 2.3).

**Text generation mode.** After establishing PROCYON's multimodal token composition capabilities as being able to process any combination of interleaved multimodal information, its abilities in text generation very clearly follow. We group both QA and phenotype generation queries under this section as both of these capabilities are fundamentally text generation; in QA, the output is a simple "yes" or "no" answer, which is supervised as such (see Extended Data Figure 2), and in phenotype generation, the output is free-form text in the form of a short description or a paragraph. After constructing multimodal inputs via the methods previously described, text generation can be done using the autoregressive mechanisms of the LLM, so no additional architectural changes are needed to enable these capabilities.

#### 2.2 Architecture specifications

The architecture of PROCYON can be broken down into several modules: the LLM, the encoders (sequence, protein structure, and ligand structure), and the connectors.

**LLM.** The base of PROCYON is a text-only, pretrained LLM. For PROCYON-SPLIT, we use LLaMA-2 (7B) [83], and LLaMA-3 [31] is used for PROCYON-FULL. This decision was made because during our development of PROCYON, LLaMA-3 models (which show far superior language modeling abilities than LLaMA-2) were publicly released as we had already began training PROCYON-SPLIT. Both LLaMA-2 and LLaMA-3 are decoder-only transformer models trained with causal language modeling objectives (see Equation 4) and can be inferenced for text generation via autoregressive generative sampling. Please reference the original publications of the LLaMA family of models [31,83] for exact specifications of the LLM's architecture.

**Protein sequence encoder.** To encode protein sequence, we use the ESM-2 (3B) model [32], which is a bidirectional, encoder-only transformer model. We freeze ESM-2 and use a mean pooling on all tokens in order to obtain one embedding vector per protein sequence. For proteins longer than the context for ESM-2 (1024 amino acids), we use a split-and-merge strategy. Thus, we split the protein into 1024 amino acid chunks, pass each chunk through the encoder in separate batches, and then pool across all output token representations for the mean pooling. This strategy ensures minimal information loss as opposed to truncation of the protein, keeping output embeddings for each amino acid and merging them in the final pooling operation.

**Protein structure encoder.** We utilized the GearNet architecture to extract embeddings from protein structures. More specifically, we chose the pre-trained variant with the multiview contrast objective, as this demonstrated superior performance across a range of tasks, based on the original results [19]. The pre-trained GearNet, sourced from the original Zenodo repository (https://zenodo.org/records/7723075), was used with its weights kept frozen throughout the ProCyon training and finetuning. We configured the GearNet model with an input dimensionality of 21, reflecting the number of input features for each node in the protein structure graph. The chosen model architecture consists of 6 hidden layers, each with 512 hidden units. Moreover, we set the edge input dimensionality to 59, and the number of relation types to 7, which corresponds to the number of relations of the relational graph network backbone model of GearNet.

**Ligand structure encoder.** We utilize the pre-trained Transformer-M model to extract embeddings from small molecule ligand structures [90], which makes use of 2D and 3D structural information of molecules. More specifically, we choose the default 12-layer ("L12.pt") configuration and weights of Transformer-M, which has been pre-trained on predicting HOMO-LUMO energy gap and 3D Position Denoising with 3.37 million by the authors [90]. The pre-trained Transformer-M was used with its weights kept frozen throughout the ProCyon training and finetuning. The 2D and 3D inputs are originally converted from SMILES and 3D coordinates, respectively, obtained through DrugBank. For molecules without 3D information, we only feed 2D inputs to the model.

**Connectors.** Connector modules bridge the gap between inputs of non-text modalities—e.g., protein sequence, protein structure, drug ligand structure—and the text tokenization space of the LLM. Each connector is a multilayer perceptron (MLP) with varying parameter sizes. The parameter size changes are a major change in architecture from ProCyon-Split to ProCyon-Full, and their exact parameter numbers are given in Supplementary Table 12 for ProCyon-Full and Supplementary Table 13. The role of these connector modules are defined in other sections. A table of all connector modules, their architecture and their respective tasks are shown in Supplementary Tables 14, 15 and 16.

**Embedding caches.** Because our encoder bases are frozen during training, we utilize embedding caches to greatly speed up training and inference. Embedding caches, which have been used in other multimodal models such as [29, 30], work by precomputing embeddings for each protein or drug ligand instance in the training dataset and stored in a matrix $\mathbf{E} \in \mathbb{R}^{N,d}$, where $N$ is the number of samples and $d$ is the dimensionality of the embedding. Thus, during forward passes, one can use an index $i$ to represent the protein sequence, protein structure, or drug structure instance and then index the embedding matrix $\mathbf{z}_i = \mathbf{E}[i, :]$. This avoids unnecessary forward passes through the frozen encoder.

#### 2.3 ProCyon inference

In this section, we describe how inference is performed on ProCyon, detailing the core capabilities. This section does not describe training objectives and optimization, which is discussed throughout Methods Sec. 3.

**Retrieval mechanism.** Retrieval inference is performed by searching over a large database of embeddings in order to find proteins most related to the input query. Given is a user input query $x_i$ and a library of amino acid sequences $\mathcal{S}$ (domains, proteins, peptides, etc.). We first compute the embedding of $x_i$ by passing it to ProCyon and then take the output embedding from the `[PROT]` token. This is then passed to the query connector $\mathbf{C}_p^q$, where a query embedding $\mathbf{z}_i$ is obtained. Then, we pass all of the amino acid sequence library $\mathcal{S}$ to the protein language model of ProCyon; this output is directly fed to the retrieval connector $\mathbf{C}_p^r$, producing an output embedding matrix $\mathbf{Z}_\mathcal{S}$. We then compute the similarity between $\mathbf{z}_i$ and the embedding matrix $\mathbf{Z}_\mathcal{S}$ (using cosine similarity) to obtain similarity scores $\mathbf{y}$. We then rank the similarity scores in descending order to obtain our ranking of similarities, and then we match the indices of this sorting with the original amino acid sequences in $\mathcal{S}$. Ultimately, we return to the user $\mathcal{S}^*$ and $\mathbf{y}^*$, which are sorted amino acid sequences and similarity scores, respectively (as is shown in Figure 1d). Users may then subset to a certain top-$k$ number of important proteins, but we leave this as a post-processing step for completeness. Such an inference is common in CLIP-style multimodal models [91], which seek to match some query to multimodal entities via a shared latent space.

This mechanism gives the user not only the ranked list of amino acids but the similarity scores as given by the model. In this way, ProCyon performs *grounded* retrieval, meaning that its outputs are not dependent on free-text generation, which is known to produce hallucinations [92], but rather specifically linked to the input amino acid library. We note that users may provide their own library of amino acid sequences or optionally use our default sequences, which are all human proteins in UniProt.

**Question answering.** At inference time for QA tasks, we use a greedy algorithm which selects the top token prediction after the "`[ANSWER]`" token provided to the model. For some tasks such as the QA filter, we use the raw logit score predicted at the "yes" and "no" tokens. However, we found empirically that a well-trained (converged) PROCYON model tended to predict almost all the softmax probability in the "yes" and "no" tokens for QA, i.e., no other tokens were included.

**Phenotype generation.** For phenotype generation (i.e., generating free-form textual responses to input instructions), we use diverse beam search [93], a modification of the beam search algorithm (a common algorithm for generating text from language models). Diverse beam search encourages separate beams to generate text distinct from the other beams, thereby resulting in semantically distinct generations; we verified this result qualitatively. In all of our experiments, we use a beam size of 10, beam group size of 2, and a diversity penalty of 0.8.

**Phenotype generation with QA filtering.** Since we generate multiple phenotype descriptions via beam search, we want to also provide some sense of confidence by the model for the generated description. We find empirically that simply ranking by log-likelihood of the sequence (as typically given by beam search algorithms) did not effectively correlate with "correctness" of the answer. For instance, the model would default to more-likely seen texts (such as more often-represented GO terms) as higher log-likelihood. Therefore, we employ a strategy called "QA filtering" where we recycle the outputs of the model and reformulate them as separate QA inputs along with the original query protein or domain. This strategy is described in Extended Data Figure 2. Then, we take the "yes" token index in the output of the PROCYON model as the certainty of the relation between the generated text and the protein/domain. This "certainty" score can be thresholded at 0.5 ($>$0.5 being "yes", $<$0.5 being "no"), or one can take the top-$k$ descriptions as ranked by the "yes" score. This requires no additional training or weights but rather an additional $N$ forward passes to the model, where $N$ is the number of generated descriptions to be filtered. This capability arises from the fact that PROCYON generates and can process in its input free-form text, even text that it generates.

QA filtering has many benefits and downfalls. The first benefit is the empirical evidence that it benefits caption quality with respect to the BERTScore metric, as seen in Extended Data Figure 3. Another benefit is that the nature of QA training gives PROCYON greater confidence on scoring more commonly-seen phenotypes, thereby providing a filter for commonly-seen descriptions to which phenotype generation might default. Let $t_i$ be a very commonly-observed textual description in one of our databases, e.g., a disease that has many associated proteins. This means that $t_i$ is seen more often by ProCyon during training, thereby giving it the chance to learn more negative examples and become more certain in predictions about this phenotype. Therefore, ProCyon is more confident in more commonly-seen examples, so if ProCyon generates a more commonly-observed text such as $t_i$, we are more confident in QA filtering to identify if this phenotype does or does not match the protein. A major downfall is that it is upper-bounded by the performance of ProCyon in QA for a given knowledge domain. If ProCyon does not converge or is not accurate for a given task or dataset description, then QA filtering is less effective. Another downfall is that seemingly-incoherent outputs may present out-of-distribution samples to ProCyon in the QA reformulation; however, this behavior was not observed empirically during testing. In fact, QA filtering was often able to catch incoherent outputs such as the middle example generated description in Extended Data Figure 2, which repeats the term "RAB9".

Overall, QA filtering provides a self-contained "check" on the outputs of ProCyon by leveraging its free-text understanding capabilities in a systematic manner to improve predictions.

**Addressing one-to-many problem through inference.** A major challenge in modeling protein phenotypes is addressing pleiotropy, the phenomenon in which a single protein is involved in multiple distinct phenotypes. Pleiotropy manifests in various ways, such as proteins that participate in multiple molecular functions, influence various diseases, or are involved in several biological pathways. For example, more than 50% of the proteins are located in multiple cellular compartments, indicating functional diversity between pathways [94], and 12% of disease-causing proteins are involved in multiple diseases [95]. This complexity breaks the assumption of a one-to-one correspondence between proteins and phenotypes, resulting in a one-to-many problem for protein-phenotype mapping.

Within ProCyon-Instruct, this complexity is evident, with proteins linked to a median of 15 phenotypes and some databases showing a mean of 21 phenotypes per protein (Gene Ontology; Supplementary Figures 13 and 14). Addressing this one-to-many protein-to-phenotype mapping is critical but remains unaddressed in existing biological foundation models. These approaches typically assume a single output $y$ for a given input $x$, thereby failing to capture the diversity of phenotypes a protein may exhibit or the multiscale nature of phenotypes.

In contrast, ProCyon addresses this challenge of complex phenotypes through both inference and training (training methods for addressing the one-to-many problem discussed in Methods

Sec. 3.1). By construction, ProCyon supports compositional queries through natural language prompts, allowing users to dynamically define phenotypes by combining multiple traits into a single, precise query – offering flexibility that stands in stark contrast to the static vocabularies employed by previous models (see Figure 2). In addition, ProCyon's inference mechanisms handle one-to-many relationships through contextual protein retrieval and conditional generation of phenotypes that capture diverse roles that proteins play in different contexts. Filtering of multiple phenotype predictions is supported by QA-filtering. Finally, the construction of retrieval outputs as ranked lists of proteins serves as a mechanism for users to explore multiple predictions as ranked by the model, allowing for inherent allowance of many proteins being predicted for a given phenotype.

##### 2.4 Implementation of ProCyon model

ProCyon is implemented in PyTorch `2.2.0+cu121` and wrapped with the Huggingface library (`transformers==4.31.0`). We follow a similar API to that of Huggingface LLMs in order to maximize user compatibility and familiarity. More details are found in our codebase, including demonstrations on how to query the model.

#### 3 Training ProCyon

In this section, we describe the training procedure for ProCyon and the design decisions made during the large-scale training of this model.

##### 3.1 Instruction tuning and interleaved multimodal inputs

To train ProCyon, we use instruction tuning, which expresses inputs in the form of an instruction, thereby increasing the model's ability to understand new, unseen tasks at inference time. Instruction tuning has been studied primarily in text-only LLMs as a way to increase performance on natural language tasks after large-scale pretraining, and it is an important alignment step in tuning large language models to more effectively follow human instructions and act as chatbots [33–37]. In this section, we will describe how to formulate instructions using interleaved multimodal inputs in order to train ProCyon.

**Instruction templates.** Because our data are not initially provided as instructions, we define templates for each task and dataset. We follow the SuperNaturalInstructions [96] formulation of instruction templates, which defines three critical components for instruction templates: 1) task definitions, 2) in-context examples, and 3) input instance.

**Task definitions.** We define task definitions as short paragraphs of text that describe the task to be performed by the model. We manually curate task definitions for each dataset across the three core capabilities of the model: QA, retrieval, and phenotype generation, as previously described. Thus, for our 20 pretraining datasets (including each relation within data sources) and three core capabilities, we have 60 task definitions. The task definition is always at the beginning of the instruction (with the exception of rephrased instructions), and it is preceded with a "`Definition:`" prefix to indicate to the language model that the following paragraph is a task definition. We release all of our task definitions along with our training data and codebase.

**In-context examples.** Following [96], we also provide in-context examples of each task to be performed by the model. In-context examples are completed, correct examples of the task to be performed. We use other (protein, description) pairs from the dataset being trained on to formulate the in-context examples. However, we fix the samples used for the in-context examples for each dataset, i.e., the in-context example given for one dataset are fixed for all example in this dataset, following the approach given by [96]. For retrieval and phenotype generation capabilities, we use one (1) in-context example, but for QA, we provide one positive example (i.e., where the answer is "yes") and one negative example (where the answer is "no"). Positive examples (including the examples for retrieval and phenotype generation) are followed by a preamble "`Positive Example:`" and negative examples are followed by a preamble "`Negative Example:`". We use these examples during inference (from the training databases) except for zero-shot tasks, where no in-context examples are provided.

Note that the ability to represent multimodal in-context examples follows directly from multimodal interleaved inputs, which allows the representation of PROCYON to be as flexible as for text-only LLMs.

**Input instance.** The input instance within the instruction is given after the task definition and in-context examples at the end of the instruction. This is the actual input query to the model, which we use for supervision during training. For text generation capabilities (QA and phenotype generation), we introduce a new token "`[ANSWER]`" which denotes to the model when it should output a response to the prompt. We train the language modeling objective (described in Methods Sec. 3.4) on the tokens given only after the `[ANSWER]` token. Full training objectives are given in Methods Sec. 3.4.

**Rephrasing instruction template components.** To increase diversity of the instructions seen during training, we use GPT-4 to rephrase the instructions given to the model. This rephrasing is in addition to the rephrasing of database descriptions that we perform for training, which is described in Methods Sec. 1.3. Here, we rephrase the task definitions that we originally write for each task. We prompt GPT-4 (version `GPT-4-1106-preview`) to generate rephrased task definitions matching the expertise levels described in Methods Sec. 1.3 and Supplementary Note 4. Our full prompt is given in Supplementary Note 5. At train time for PROCYON-FULL, we sample first the expertise level and then the rephrasing level. Expertise level is tied across both task definitions and entity descriptions, but rephrasings are sampled independently for each of these components. This means that the task definition and the input description given for an example will be from the same expertise level, but these may be given as a summary and a rephrasing, a rephrasing and a simplification, etc. We then use a "personality system prompt", as shown in Figure 4e, which tells PROCYON to process inputs and generate outputs that match the style of the requested expertise level. These prompts are given in full within Supplementary Note 4.

**From protein-text pairs to each PROCYON capability.** PROCYON is trained for the three capabilities discussed in Methods Sec. 2.1. For enabling these capabilities, we design instructions that match each capability. Each sample from our databases (excluding the interaction knowledge domain which is discussed in the subsequent section) is a protein, text pair $(p, t)$, where $p$ is associated to the phenotype described by the textual description $t$. Let $\mathcal{F}$ denote the PROCYON model and $I$ denote some human-provided instruction (encoded via the instruction tuning formatting). Then, each task can be broken down as a mapping between $p$ and $t$ conditional on instructions. Define each mode of PROCYON with $\mathcal{F}^{\text{R}}$ as the retrieval mode, $\mathcal{F}^{\text{QA}}$ as the QA mode, and $\mathcal{F}^{\text{C}}$ as the phenotype generation mode, then each function is defined as such:

$$\mathcal{F}^{\text{R}}(t|I) \mapsto p; \mathcal{F}^{\text{QA}}(p, t|I) \mapsto \{\text{“yes”}, \text{“no”}\}; \mathcal{F}^{\text{C}}(p|I) \mapsto t \quad (1)$$

Thus, we construct instructions to reflect these tasks. For retrieval instructions, we give the model the instruction and text, prompting for the protein that matches the query text and instruction combination. For QA instructions, we give the model the instruction, protein, and text and prompt the model to predict if the text describes a property of the protein ("yes") or not ("no"). For phenotype generation instructions, we give the model the instruction and protein, prompting the model to generate the text $t$ that matches the protein. In this way, one pair of protein-text $(p, t)$ is transformed into three samples phrased in different ways to the model via instruction tuning.

**Instructions for molecular interactions.** Molecular interactions, including protein-protein interactions for the STRING dataset and protein-peptide interactions for the PDB dataset, are encoded as instructions. As a reminder, PROCYON can handle interleaved multimodal inputs (see Methods Sec. 2.1), so encoding protein-protein interactions is straightforward. Denote a protein-protein (or peptide-protein for generality) sample as $(p_i, p_j, r)$, where $p_i$ and $p_j$ are two proteins and $r$ is the relation describing their relationship, such as homology, coexpression, or experimental interaction. Without loss of generality, this can also refer to a general physical interaction such as binding that is referred to in the PDB dataset. We encode the relation as part of the instruction, which we can denote as $I(r)$ for the relation-augmented instruction. This is as simple as mentioning to the model that the relation being predicted is, for example, coexpression. Then, we can construct retrieval and QA examples as:

$$\mathcal{F}^{\text{R}}(p_i|I(r)) \mapsto p_j; \mathcal{F}^{\text{R}}((p_i, p_j)|I(r)) \mapsto \{\text{“yes”}, \text{“no”}\} \tag{2}$$

Thus, we train PROCYON models on retrieval and QA examples from datasets in the interaction knowledge domain. Phenotype generation is omitted because no text is naturally associated with this interaction data. All instruction templates, including task definitions, used in our training process is available in our codebase which can be found at https://github.com/mims-harvard/ProCyon. Example instructions are shown in Extended Data Figure 5, with each component highlighted that is discussed in this section.

#### 3.2 Context augmentation

We introduce several strategies to improve model training through methods known as context augmentation (CA). Because input to PROCYON can be expressed flexibly as interleaved multimodal-text data, the model can encode arbitary context to perform a task. Therefore, we design strategies that leverage domain-specific intuition to increase PROCYON's performance. We define three kinds of CA techniques, Ontology CA, Disease CA, and Drug CA.

Ontology CA utilizes the ontologies of Gene Ontology and Reactome databases to retrieve upstream ancestors in the ontology to insert as context to the input prompt; this narrows the search space for PROCYON given the high number of protein-phenotype connections. Disease CA retrieves a functional description of the protein during training to further connect the molecular phenotypes to high-level clinical descriptions of diseases. Drug CA uses the indication and mechanism of action to boost prediction of other given properties of the molecule that are relevant to therapeutics tasks. More details are given in Supplementary Note 24, 25 and 26 and in Supplementary Table 17. A summary of all datasets that use each CA technique is given in Supplementary Table 18.

#### 3.3 Data processing details for training PROCYON

In this section, we summarize a collection of design choices not outlined in previous sections that were helpful for training PROCYON.

**Epoch scaling per task.** We implement an epoch scaling technique to combat overfitting on textual data during training of PROCYON. Since we train PROCYON to simultaneously perform retrieval, QA, and phenotype generation tasks, we must carefully balance the training resources used for both of these main tasks. To balance such differences during training, we scale the frequency of training for the main tasks (QA, retrieval, and phenotype generation). Define a standard whole-task training batch at step $i$ as $\mathbf{B}_i = \{B^i_{\text{QA}}, B^j_{\text{R}}, B^k_{\text{C}}\}$ where $B^i_{\text{QA}}, B^j_{\text{R}}, B^k_{\text{C}}$ are batches of QA at QA step $i$, retrieval at retrieval step $j$, and phenotype generation (captioning) at captioning step $k$, respectively. An epoch scaling factor is an integer $n_\tau$ associated to one of the given tasks, thus $n_{\text{QA}}, n_{\text{R}}, n_{\text{C}}$ is the epoch scaling factor for QA, retrieval, and captioning, respectively. $n_\tau$ controls the frequency at which a task is seen during a given training step; for example, a set of $n_{\text{QA}} = 2, n_{\text{R}} = 1, n_{\text{C}} = 1$ would yield a training step as $\{B^i_{\text{QA}}, B^{i+1}_{\text{QA}}, B^j_{\text{R}}, B^k_{\text{C}}\}$. Over training, this amounts to each task being seen at different numbers of epochs. Epoch scaling parameters for PROCYON are reported in Supplementary Table 19.

**Two-stage training.** For training PROCYON-SPLIT, we utilize a two-stage training process whereby the connector layers are pretrained using limited data while keeping the language model frozen. This was originally inspired by [84], which follows a similar architecture but for vision-language models. In this work, they pretrain the connector layers while keeping the language model frozen, thereby providing a better initialization of connector weights before tuning language model weights in the second stage. Thus, we trained a model PROCYON-STAGE1 on the UniProt dataset without using instruction tuning. This means that the model is prompted with a simple retrieval query or caption query without the added text of an instruction. This allows the newly-introduced tokens to be tuned as well as the connector layers. We report PROCYON-STAGE1 hyperparameters alongside hyperparameters for other versions of ProCyon in Supplementary Tables 19 and 20. More details on the dataset used to train ProCyon-Stage1 is included in Supplementary Note 27.

For training ProCyon-Full, we decided to not follow this protocol since work that was published during the duration of training ProCyon-Split suggested that the two-stage training procedure is not necessary [97].

**Integrating rephrasings during training.** During training of ProCyon-Full, we integrate rephrasings for task descriptions (Methods Sec. 3.1) and phenotypes as a sampling procedure. For each sample drawn in a minibatch, we decide with 0.5 probability if that entity will be rephrased, for the entities that we consider to be rephrased (see Supplementary Table 11). This probability was determined via internal testing on a limited model that is not reported here. After deciding that an entity will be rephrased, we then select the expertise level and granularity level independently and uniformly. Please see Methods Sec. 3.1 for more information on how the rephrased instructions are constructed.

#### 3.4 ProCyon training objectives

We formalize ProCyon training into three primary task types (and their shorthand for this section): protein retrieval and prioritization ("retrieval"), question answering ("QA"), and phenotype generation ("captioning"). In this section, we describe the training procedures and objectives for each of these tasks.

**Retrieval training.** We train the model for retrieval based on contrastive learning, which optimizes a joint embedding space for outputs from the protein language model and from the LLM. Specifically, we adopt the InfoNCE [98] framework for multimodal representational alignment, which has been a popular choice in vision-language models [91] as well as some protein-language models [29]. This is formalized as an output text embedding $\mathbf{z}_i^t$ and output protein embedding $\mathbf{z}_i^p$. Thus, our objective is:

$$\mathcal{L}_{\text{ret}}(\theta) = \frac{1}{N}\sum_{i=1}^{N} \log \frac{\exp(\text{sim}(\mathbf{z}_i^t, \mathbf{z}_i^p)/\tau)}{\sum_{j=1}^{N} \exp(\text{sim}(\mathbf{z}_i^t, \mathbf{z}_j^p)/\tau) \cdot \mathbf{m}_{i,j}} + \frac{1}{N}\sum_{i=1}^{N} \log \frac{\exp(\text{sim}(\mathbf{z}_i^p, \mathbf{z}_i^t)/\tau)}{\sum_{j=1}^{N} \exp(\text{sim}(\mathbf{z}_i^p, \mathbf{z}_j^t)/\tau) \cdot \mathbf{m}_{i,j}} \tag{3}$$

Where $\theta$ are the weights of ProCyon, $N$ is the batch size, $\mathbf{z}_i^t$ is the retrieval embedding from the LLM, $\mathbf{z}_i^p$ is the retrieval embedding from the PLM, $\tau$ is a learnable temperature parameter, and

$\mathbf{m}_{i,j}$ is a mask between samples $i, j$ to account for mismatches in negatives across the batch (i.e., batch conflicts due to in-batch negative sampling). Note that the denominator in either term is guaranteed to be non-zero since by definition, $\mathbf{m}_{i,i} = 1$.

**Sampling negatives for Retrieval Optimization.** We perform in-batch negative sampling, as is standard practice in contrastive learning frameworks due to efficiency and simplicity [99, 100]. In the distributed training framework, we pool negative samples from all proteins across the global batch for each forward pass; such an operation is accomplished through an `all_gather` call which is implemented in PyTorch distributed [101]. This effectively allows us to increase our batch size to leverage all protein embeddings across the global training of the model. However, we found that during development, the model encountered many false-negatives when using naive in-batch negative sampling, leading to signficant conflict in retrieval training. Therefore, we also implement a negative sampling mask, whereby when a protein is falsely sampled as an in-batch negative sample, we set the distance to 0 in the distance matrix for the conflicting pair. Further details can be found in the PROCYON codebase.

**Text generation training.** For text generative tasks, we use causal langauge modeling as an objective. As previously discussed, we align QA and phenotype generation tasks (as discussed in Figure 1) under "text generation" for the purposes of training. Both QA and captioning are trained as language modeling tasks, using the causal language modeling objective. Let a training sample $X$ be broken down into a sequence of tokens $\{x_1, x_2, \dots, x_N\}$. For our purposes, we mask out the part of the sequence that consists of the instruction, so let the instruction be tokens $\{x_1, x_2, \dots, x_M\} \subset X$ where $M < N$. Thus, our causal generative loss is defined as such:

$$\mathcal{L}_{\text{gen}}(\theta) = -\sum_{i=M}^{N} \log P(x_i \mid x_1, x_2, \dots, x_{i-1}; \theta) \tag{4}$$

Both QA and captioning tasks are trained on this objective. Please consult Methods Sec. 3.1 for more information on the content of text trained on within the instructions.

**Loss rescaling by dataset.** Each dataset in our training set contains different numbers of samples, meaning that there is high imbalance between tasks during training. For example, function prediction via Gene Ontology makes up about 29.6% of the dataset, while drug target prediction via DrugBank only accounts for 2.1% of the samples. Further, our datasets are highly non-injective, meaning that one text may be overrepresented for many proteins. For example, following the previous example, Gene Ontology datasets have 52.2 proteins per textual description, e.g., a given text is seen 52.2 times on average per epoch during training; in contrast, drug target prediction via DrugBank has 4.5 proteins per textual description. This leads to the possibility of the model overfitting on descriptions from Gene Ontology and other databases with high levels of this non-injectivity. To combat this task imbalance and the non-injectivity of the datasets, we implement a manual loss rescaling method whereby the loss from rarely-seen databases are upweighted for open-ended phenotype generation. The choice of rescaling weights was made based on a combination of task imbalance as well as observed difficulty on tasks learned by training ProCyon-Split.

Practically, loss rescaling is implemented as a constant multiplier $\lambda_{\mathcal{D}_i}$ for each dataset $\mathcal{D}_i$. Formally, for some loss function $\mathcal{L}$ applied on example $\mathbf{x}$ for a batch on dataset $\mathcal{D}_i$, the loss-rescaled version $\mathcal{L}^s$ is defined as:

$$\mathcal{L}^s(\mathbf{x}|\mathcal{D}_i) = \lambda_{\mathcal{D}_i}\mathcal{L}(\mathbf{x}) \tag{5}$$

We implement loss rescaling on ProCyon-Full but not ProCyon-Split, as this phenomenon was observed while training ProCyon-Split. Values used for the loss rescaling are shown in Supplementary Table 21.

**Total loss function.** In our final loss function, we perform a sum across the QA, retrieval, and captioning batch losses. Assume we receive a batch $B_{\text{QA}}, B_{\text{R}}, B_{\text{C}}$; note that with epoch scaling (discussed in Methods Sec. 3.3), not every batch may be constructed this way, but consider this as a batch WLOG. Consider each batch to come from datasets $\mathcal{D}_{B_{\text{QA}}}, \mathcal{D}_{B_{\text{R}}}, \mathcal{D}_{B_{\text{C}}}$, which are the datasets for the QA, retrieval, and captioning batches, respectively. Let $\mathcal{L}(B)$ denote the loss function computed over a batch $B$, i.e., over all samples in the batch. Thus, our total loss function $\mathcal{L}_{\text{ProCyon}}$ is defined as such:

$$\mathcal{L}_{\text{ProCyon}}(B_{\text{QA}}, B_{\text{R}}, B_{\text{C}}) = \lambda_{\text{QA}}\mathcal{L}^s_{\text{gen}}(B_{\text{QA}}|\mathcal{D}_{B_{\text{QA}}}) + \lambda_{\text{R}}\mathcal{L}^s_{\text{ret}}(B_{\text{R}}|\mathcal{D}_{B_{\text{R}}}) + \lambda_{\text{C}}\mathcal{L}^s_{\text{gen}}(B_{\text{C}}|\mathcal{D}_{B_{\text{C}}}) \tag{6}$$

Where $\lambda_{\text{QA}}, \lambda_{\text{R}}, \lambda_{\text{C}}$ correspond to scalar weights for the QA, retrieval, and captioning tasks, respectively. We then optimize our model for this loss function with the AdamW optimizer.

**Targeted modules during training.** During training, we optimize the weights of the LLM and the weights of the connectors. The weights of the protein language model (PLM), protein structure encoder, and ligand structure encoder are frozen and not tuned. These decisions follow results from popular vision-language models such as LLaVA [84] which, in a similar architectural setup, train the LLM and connector (analagous to our connector modules) with full-finetuning while freezing the vision encoder (to which our multimodal encoders are analagous).

##### 3.5 Ablations for PROCYON

To better understand the crucial components of PROCYON, we conduct a number of ablations. Our ablations are **inference-time ablations**, whereby we seek to understand which parts of the inference mechanisms are important for PROCYON'S performance. This is to better understand user recommendations for querying PROCYON. Please see limitations below for a discussion on this point. All ablations are made on PROCYON-SPLIT in order to evaluate fairly on held-out datasets.

**Ablation 1: Structure token.** We ablate the use of the structural token for QA prediction tasks. QA is the sole focus for this experiment because structure is not explicitly used in the retrieval task. PROCYON shows little-to-no change in performance when removing the structural token from the input, demonstrating the robustness of our model when protein structures might be poorly-characterized or not experimentally-determined (Supplementary Table 22).

**Ablation 2: ICL examples.** In this experiment, we consider the case where in-context learning (ICL) examples are excluded from the input. ICL examples are intrinsically a part of the instruction tuning framework used to train PROCYON (Methods Sec. 3.1). We find that taking out ICL examples has approximately no effect for retrieval (Supplementary Table 23) while for QA, there seems to be a negligible negative impact on performance when excluding ICL examples compared to their inclusion (Supplementary Table 24). We hypothesize this is due to the ICL examples providing a template, which is important for narrowing the options of the LLM when answering QA-style questions, i.e., limiting answers to "yes" or "no". Thus, we recommend using an ICL example for QA tasks with PROCYON, but for retrieval, no ICL examples are needed.

**Ablation 3: Context augmentation.** In this ablation, we take out the context augmentation (Methods Sec. 3.2) at inference time to understand if the model uses the CA mechanism. We ablate three CA mechanisms: ontology CA , disease CA , and drug CA. For ontology CA, we see no significant drop in performance when removing CA from the input prompt; in fact, performance slightly improves when removing CA, most likely due to the stochasticity of choosing upstream neighbors at inference time (Supplementary Table 25). For disease CA, we see a slight drop in performance in DisGeNet and more mixed results in OMIM (Supplementary Table 26). Because DisGeNet is far larger than OMIM, we consider these results to be more reliable, thus we can conclude the disease CA provides a marginal benefit to modeling performance. For drug CA, we clearly see that it is crucial to model performance, with performance dropping to almost random (about $0.5$ accuracy) when omitting it (Supplementary Table 27). This is most likely due to the challenging nature of drug prediction tasks, which require extensive knowledge of biological systems as well as molecular components of both drugs and proteins. Thus, the use of drug CA demonstrates how the free-text capabilities of PROCYON can be used to produce more compelling results, a property that is not achievable by many baseline models compared in this study.

**Limitations.** Because of the high cost of training PROCYON, we do not conduct training ablations, i.e., we do not train separate instances of PROCYON to understand training choices. We acknowledge that this is a limitation of our work; however, we note that the monetary and energy cost is prohibitive given the long training times for our model (see Supplementary Table 28). We plan for future work to explore the design components of the PROCYON architecture and training procedure. We hope that as the community explores similar architectures to that of PROCYON, we can derive a better understanding of the design decisions for training such a model.

#### 3.6 PROCYON training implementation

**Training duration.** Training PROCYON was a significant effort that included large numbers of GPUs and many days of training on a large distributed cluster. Training duration can be measured in either a data-centric manner or a compute-centric manner. By a data-centric method, we can measure train duration in numbers of epochs, where an epoch is one iteration over every sample for all training datasets considered in that training. We also break down our reporting of training epochs by task, e.g., QA, retrieval, and caption. Exact training epochs are found in Supplementary Table 29.

Using a compute-centric method to calculate training time, we measure training time in terms of wall clock duration of training iterations. GPU time is then calculated by multiplying the wall clock duration of training with the number of GPUs used during training. Note that this does not correspond to raw GPU usage, as some CPU time may be considered during the training iterations for non-GPU operations such as instruction construction. The total GPU training time for ProCyon-Full was 287.28 days, ProCyon-Split was 173.91 days, and ProCyon-Bind was 2.39 days. Full train time statistics are found in Supplementary Table 28.

**Training hardware.** We train all versions of ProCyon on Nvidia H100 GPUs provided by the Kempner Institute for the Study of Natural and Artificial Intelligence at Harvard University. We used 48 GPUs to train ProCyon-Split and 32 GPUs to train ProCyon-Full; this was due to shifting resource availability, explaining why ProCyon-Full resulted in longer training time overall.

**Training infrastructure.** To train ProCyon, we use state-of-the-art methods in distributed neural network training. For our data and model-parallelism, we use Sharded Data Parallel (ZeRO Stage 2) as implemented in the Deepspeed library [102] version `0.12.4`. ZeRO Stage 2 conveniently keeps the data parallel world size equal to the number of GPUs being used, so each GPU was used for parallel data iteration, as implemented by Deepspeed. More details on our full training stack are available in our codebase (https://github.com/mims-harvard/ProCyon).

##### 3.7 Addressing one-to-many problem through training

By training on multiple knowledge domains, ProCyon achieves multiscale phenotypic resolution, enabling precise querying across molecular functions, diseases, pathways, and therapeutics. Further, context augmentation integrates additional information about phenotypes into the model, significantly improving predictive performance. By training on natural language descriptions, this allows for flexible definition of phenotypes, which can capture complexities beyond coarse-grained annotations that map to many proteins (Supplementary Figures 13 and 14).

#### 4 Analysis and application of ProCyon

We now describe the methods used for various analyses, benchmarking, and probing of ProCyon conducted throughout this work.

##### 4.1 Benchmarking and evaluation

When benchmarking question answering and retrieval performance, we wanted to ensure fair comparison of ProCyon to our various baselines. For this purpose, we implemented a standardized evaluation framework and integrated all baselines within this framework. The desired datasets and splits are specified via a YAML config file, and similarly the models to be benchmarked are specified in a separate YAML config file alongside any relevant model-specific arguments. Each model is wrapped in a standardized API, allowing us to use the same data loading and evaluation logic across all models to ensure they're evaluated fairly and on the same set of examples.

All benchmarking of PROCYON was performed using the PROCYON-SPLIT model, and all benchmarking results use evaluation splits only, averaged across non-zero shot splits (i.e. many-shot and few shot) unless explicitly specified as zero-shot results.

**Question answering evaluation.** For each dataset and split, question-answer pairs are constructed in batches. Each batch consists of eight positive (protein, text) relationships, which are then supplemented with eight negative (protein, text) relationships. Negative relationships are constructed as follows: for each positive (protein, text) pair, a single protein is sampled from the set of all proteins in our overall dataset, with sampling probability according to dissimilarity to the selected protein (i.e. biasing towards less similar proteins). The sampled protein and the original text then form a negative example. In this manner, each batch contains a balance of positive and negative examples for QA evaluation.

Each batch of (protein, text) pairs is then passed to a given model following a standardized API, and the model returns a single binary prediction for each received example. Accuracy is calculated as the mean number of correct predictions across the entire dataset split. 95% confidence intervals for accuracy are calculated via bootstrapping using scipy's (v1.13.0) `bootstrap` function: `bootstrap(n_resamples=9999, method='BCa', confidence_level=0.95)`.

**Retrieval evaluation.** For each dataset and split, we first identify the set of (protein, text) relationships to evaluate. For a given text, we begin by treating all proteins it's related to in this split as the positive retrieval targets, and all other proteins as negative targets. To avoid conflict with training data, we remove all (protein, text) relations in the training set from consideration. For example, this avoids treating a (protein, text) relation in the train set as a negative relation at test time. This is necessary because our splits are constructed at the level of (protein, text) relations, and thus a given text can appear in multiple data splits. In this manner, we construct a cleaned set of positive and negative retrieval targets per text.

The set of texts and the full set of proteins in our overall datasets are then passed to a given model following a standardized API, and the model returns a score for each (protein, text) pair, where a higher score corresponds to higher retrieval ranking.

We evaluated each model's retrieval performance using $F_{\max}$, the maximum $F_1$ score achieved across all possible thresholds for separating positives and negatives. To evaluate a model's performance across the entire dataset split, we first calculate $F_{\max}$ for each text separately, using only its positive protein relations and considering all other proteins as negatives. To account for severe label imbalance (most texts relate to very few of our $\approx$18,000 proteins), we randomly sample multiple negatives per positive sample. In particular, for each text, we sample 10 negative proteins for each positive, or if this number would exceed the total number of proteins, we use the full set of proteins as negatives. We perform this sampling 5 times and return the average of the samples as the $F_{\max}$ score for this text. The final $F_{\max}$ score for the entire dataset split is then the average of the $F_{\max}$ scores of each text. Similarly to QA, 95% confidence intervals are calculated via bootstrapping at the text level (i.e. resampling per-text $F_{\max}$ scores) using scipy's (v1.13.0) `bootstrap` function: `bootstrap(n_resamples=9999, method='BCa', confidence_level=0.95)`.

#### 4.2 Baseline model methods

We compare PROCYON against a number of baseline models to evaluate it's capabilities with respect to different types of modeling approaches. This includes five protein-only models, including sequence-only models BLAST [38] (kNN approach), GearNet [19] (both kNN and MLP approach), and ESM-2 [32] (both kNN and MLP approach). We also compare to multimodal models, including ProtST [29], BioTranslator [28], ProtLLM [30], and ESM-3 [16] (both kNN and MLP approach). Detailed information on the implementation of these baseline models is included in Supplementary Notes 9 and 10.

#### 4.3 PROCYON retrieval performance on varying evidence sources

These results are shown in Figure 3b. To validate the abilities of PROCYON to retrieve proteins across different diseases using as ground truth (disease, gene) associations derived from OpenTargets (OT) [66] and collected through different evidence sources (e.g., genetic associations, somatic mutations, animal models, affected pathway, literature and known drug). For each disease, we derive the direct disease-gene associations for each data source, resulting in 3372 diseases being included in the analysis. Exact details on data curation are included in Supplementary Note 12.

We query PROCYON using the following task description:
"You will be shown text describing a disease, which could be common, rare (Mendelian), complex or environmental. Your job is to determine the proteins that are associated with the disease described. The associations are provided by various expert curated resources, representing relationships where there is established evidence linking genetic variations or mutations to the occurrence,

risk, or progression of diseases. The associations indicate whether a protein, through its variation or mutation, plays a role in the manifestation of a disease, either by directly causing it, serving as a marker for it, being implicated in its mechanism, or being targeted for therapeutic intervention."

We use the disease description available in OMIM as input description, and retrieve the ranked list of proteins. Finally, for each disease and source, we derive the ranked percentile for each protein in the ground truth list. The retrieval percentile is a metric used throughout retrieval evaluations, which refers to the percentile of the rank of the ground-truth protein with respect to all human proteins considered in our model. In this instance, we calculate the median over the set of all proteins in the ground-truth list for each disease, resulting in a median percentile score per disease and source.

#### 4.4 Disease gene prioritization with increasing evidence

This section addresses the experiment discussed in Figure 3c. The list of diseases and input prompts are derived from the Diagnostic Manual of Mental Disorders (DSM) fifth edition. The DSM provides the standard classification of mental disorders that is commonly used by mental health specialists in the United States. It provides a structured text that is characterized by multiple sections that are consistent across the majority of the diseases. For this analysis, we used only the sections "Diagnostic Features" and "Associated Features supporting Diagnosis" [41].

To select the diseases to query the model for, we first identify the subset of diseases available both in the DSM-5 and in OpenTargets (OT). We complete the mapping from DSM-5 to OT manually by using the OT online dashboard to check for diseases https://platform.opentargets.org/, resulting in 63 diseases being included in the analysis. Then, when available, we manually extract the "Diagnostics" and "Associated Features supporting Diagnosis" descriptions for each disease, resulting in 51 diseases.

To assess the performance of ProCyon, we extract the overall ground truth direct gene-disease associations from OT (data were downloaded on the 10th June 2024) for the 51 diseases, regardless of the database and source used to derive them. We apply a threshold of 0.8 on the gene-disease associations confidence scores, in order to keep only high confidence associations. We conduct the analysis only on diseases with at least one gene-disease association that passed the threshold, resulting in 30 diseases.

Finally, we query ProCyon with a task definition that reads: "You will be shown text describing a disease, which could be common, rare (Mendelian), complex or environmental. Your job is to determine the proteins that are associated with the disease described. The associations are provided by various expert curated resources, representing relationships where there is established evidence linking genetic variations or mutations to the occurrence, risk, or progression of diseases. The associations indicate whether a protein, through its variation or mutation, plays a role in the manifestation of a disease, either by directly causing it, serving as a marker for it, being implicated in its mechanism, or being targeted for therapeutic intervention." We query the model five times for each disease, each time using a different input description. We first query the model just with the disease name, the "Diagnostics Features" and the "Associated Features supporting diagnosis". Then, to increase the amount of evidence input into the model, we query ProCyon using "Disease + Diagnostics Features", and "Disease + Diagnostics Features + Associated Features supporting diagnosis". Finally, for each disease, we derive the ranked percentile for each protein in the ground truth list and calculate the median, resulting in a median percentile score per disease.

#### 4.5 Retrieving proteins for composite and complex phenotypes

**Retrieving pleiotropic proteins.** This addresses the result in Figure 3e. For this experiment, we consider pleiotropic proteins as proteins annotated to more than one GO biological process terms that are not an ancestor of one other and do not share any offspring GO terms ("non-overlapping"). With this, we consider GO biological process terms above level 2, and curate mutually non-overlapping pairs, triplets, and quadruplets of terms that have between 3 to 20 overlapping protein annotations. Due to the vast combinatorial space, we restrict our experiment to randomly sampled 1000 pairs, 500 triplets, and 200 quadruplets (decreasing number because of permutation of queries required as specified below). For each pair, triplet, or quadruplet of signaling pathways, we query ProCyon in different ways and obtain the quantiles of the associated pleiotropic proteins.

In the "composite query" approach, we query ProCyon with a task definition defining the retrieval target, along with the descriptions of pathways for each pair, triplet, or quadruplet of pathways. Specifically, the task definition reads: "You will be shown text descriptions of two[three/four] biological processes. Your job is to determine if a protein is involved in both processes described. Involvement is determined by whether or not there is evidence from an experiment directly supporting the annotation of the protein." For each pair, triplet or quadruplet of pathways, we query the model with all permutations of pathway description combinations (2 for pairs, 6 for triplets and 24 for quadruplets). The quantiles of proteins from each query are aggregated with geometric mean.

In the "aggregated individual ranks" approach, we query PROCYON with a task definition defining the retrieval target, along with the description of one of the two, three, or four pathways taken from its mapped GO biological process term. Specifically, we adopt the same task definition as used for the biological process of GO. For each pair, triplet or quadruplet of signaling pathways, we query the model twice, three times, or four times, each time with a different pathway. The quantiles of proteins from each query are aggregated with geometric mean.

In the "random composite query" approach, we query PROCYON with a task definition defining the retrieval target, along with the descriptions of two, three, or four randomly sampled GO biological process terms from our database. The task definition is identical to the "composite query" case. For each pair of pathways, we query the model with all permutations of pathway description combinations. The quantiles of proteins from each query are aggregated with geometric mean.

We also compare PROCYON on this task to ProtST, one of the leading baseline methods that utilizes free-text inputs. To remain consistent, we input the same description to ProtST and use the same ranking mechanisms.

**Retrieving STING for neuronal inflammatory stress response.** This point addresses the result in Figure 2h. Our latest latest accession date of any database used for training is Oct 2023 (GtoP), nine months before the publication of [42] in July 2024. Using the queries in Supplementary Note 14, we query the model with increasing information about the STING pathway to observe it's ability to prioritize scientific evidence in retrieving proteins. Each level of text is permuted 100 times by randomly dropping 5% of words to obtain confidence intervals (shown as error bars). All queries are subsequently fed into PROCYON to obtain the quantiles of STING along with a confidence interval on the bootstrap procedure. For embedding space generation in Figure 2h, we use the same methods as described in Supplemental Note 1 for the protein embeddings. For prompt embeddings, we superimpose the embeddings onto the protein embedding space by taking the mean embedding of the top-5 most similar proteins to each prompt, as defined by ProCyon. This is done to visually align the embedding spaces, where spatial proximity is lost in the UMAP projection.

#### 4.6 Comparisons to text-only LLMs

This section addresses the comparison to text-only LLMs in Figure 3. To understand how adding rich and generalizable protein representations enables ProCyon to reason over proteins, we compared our model against text-only LLMs. For this evaluation, we focused on the question answering and phenotype generation tasks, and compared to state-of-the-art LLMs at a range of model sizes.

For a fair comparison, we prompt all LLMs with the same prompts used for ProCyon, but with differing methods of how the protein is encoded in text. Similarly to ensure consistency with ProCyon evaluation, we provide an example of the desired output style in the prompt to the LLMs. We probe the ability of text-only LLMs to understand proteins across a range of representations by considering three possible text encodings of proteins: amino acid sequence (-AA), UniProt ID (-UP), and HGNC gene ID (-HG). In each case, we directly replace ProCyon's protein sequence and structure tokens with the corresponding encoding. We acknowledge that the amino acid representation is likely to perform very poorly due to full protein amino acid sequences being unlikely to appear in the training corpuses of text-only LLMs; however, we also note that this is the only representation that can accommodate completely novel or unseen proteins, as the UniProt and HGNC encodings are limited to proteins which have received such IDs.

For our evaluations, we consider LLMs across three different size classes. First, models with parameter count similar to ProCyon (∼1X size of ProCyon): Meta-Llama-3-8B[2] and Meta-Llama-3-8B-Instruct[3]. Both Llama models were queried using the `transformers` Python package, with text generation parameters: `max_new_tokens=300, top_p=0.9, temperature=0.7`. For models with parameter count larger than ProCyon (∼10X size of ProCyon): Mistral Large - queried via Microsoft Azure AI Services Chat Completions API. API version `2024-05-01-preview`, model `mistral-large`, temperature of 0.5, and max tokens of 350. For models with parameter count significantly larger than ProCyon (>50X size of ProCyon): GPT-4o - queried via Microsoft Azure AI Services Chat Completions API. API version 2024-05-01-preview, model `gpt-4o`, temperature of 0.5, and max tokens of 350.

For all LLM comparisons, we use the ProCyon-Full model to ensure that our model has exposure to as much data as possible for a better comparison to the internet-scale training corpuses of the LLMs we compare against.

[2]https://huggingface.co/meta-llama/Meta-Llama-3-8B

[3]https://huggingface.co/meta-llama/Meta-Llama-3-8B-Instruct

**QA evaluation.** Due to the scale of our datasets, we have an extremely large number of QA prompts to evaluate over, even when only considering positive QA examples, i.e., question-answer pairs generated from the relations in our dataset, where the ground truth label would be positive. Since evaluating all such QA prompts would be prohibitively expensive, we subsample our datasets to maintain a reasonable cost. For each dataset and split, we randomly sample 128 question-answer pairs, where the sampled set consists of 64 positive examples and 64 negative examples. Since we have 3-4 splits per dataset, this corresponds to roughly 400-500 prompts per dataset, which strikes a reasonable balance between sample size and cost.

Given the selected prompts, we generate QA results for the selected LLMs by using the text-only prompts with proteins encoded using one of the methods described above. Our prompt explicitly states "You may only output yes or no", to which PROCYON and GPT-4o adhere, but other models require more post-processing. In particular, we found that Mistral Large tended to output rationales before outputting an answer in the format "Output: <answer>", whereas the Llama-3-8B models often didn't output an answer at all. To extract outputs from LLM answers, we first check if the answer begins with "[Yy]es" or "[Nn]o" and use that as the answer if so, otherwise we use the regex "`Output: (\w+)`" to attempt to recover a yes or no answer. We find that this approach identifies a yes or no answer for all PROCYON responses, all GPT-4o responses, and all but 2 Mistral Large responses, with Llama-3-8B models frequently outputting nonsensical text.

Given the extracted yes or no answers from each LLM, we are then able to proceed with QA evaluation as performed for our baseline models, including bootstrapping for 95% confidence interval calculation. For Llama-3-8B models in particular, we treat outputs without a yes or no answer as incorrect, resulting in binary accuracies below 50%.

**Phenotype generation evaluation.** Similar to QA, we have to downsample our datasets for evaluating text-only LLM's phenotype generation performance. To ensure we sample proteins with a range of representation in scientific literature (and by proxy, representation in an internet-scale LLM training corpus), we use UniProt annotation scores as a proxy for how well-studied a given protein is. UniProt annotation scores range from 1 (least studied) to 5 (most studied). We select three of our datasets for evaluation: UniProt, Reactome, and DisGeNet, which each represent distinct knowledge domains. Within each dataset, we select all proteins with annotation scores of 1 or 2, and randomly sample up to 50 proteins each per annotation score of 3, 4, and 5, resulting in a total of 657 proteins across all three datasets.

For phenotype generation from ProCyon, we use Diverse Beam Search [93] with 5 beam groups each of size 2, resulting in a total of 5 generated phenotypes per prompt. We then select amongst these captions using our QA filtering process, in which each caption is passed back to the model alongside the corresponding protein as a question answering prompt. The resulting probability of the "yes" token is then used as a confidence metric for the caption, and for each protein, we retain the single generated phenotype with the highest confidence.

Text generation for text-only LLMs is performed using the same prompt as is used for ProCyon, except with the protein token replaced with the corresponding textual representation of the protein. Text generation parameters are as listed above.

Given a set of generated phenotypes for a single protein, we generate reference-based performance metrics compared to the set of ground-truth annotation texts from the corresponding database. As a single protein may be associated with multiple phenotypes (e.g., a target of multiple drugs, or association with multiple diseases), a protein can have multiple reference phenotypes to compare against. To capture the case in which the generated text matches one association, but none of the others, we calculate metrics against all reference texts, and take the max of these values as the single metric by which to evaluate the generated text. For the performance metric, we use BERTScore F1 [43], an extension of n-gram matching metrics that also accounts for semantic similarity between words rather than requiring exact matches (e.g. the words "support" and "maintain" can be a high-scoring match). We calculate BERTScore with the parameters `rescale_with_baseline=True, lang="en-sci"`, and the hashcode `allenai/scibert_scivocab_uncased_L8_no-idf_version=0.3.12(hug_trans=4.31.0)-rescaled`. Given a single BERTScore F1 metric for each protein in the evaluated dataset split, we then calculate 95% confidence intervals via bootstrapping as previously described.

**LLM-based holistic evaluation.** To perform LLM-based ranking, we provide the judge LLM with the original phenotype generation prompt, the generated text from ProCyon, the generated text from another model, and the set of reference texts for the given protein. For the ProCyon outputs, we select the top caption given by the diverse beam search generation algorithm. In addition, we insert a brief preamble to attempt to match the distribution of chat-based LLMs. The preamble states: "Here's a description for the protein <protein_id>", where "<protein_id>" denotes the ID of the protein being examined. We prompt the model to select which of the two models' outputs is more correct relative to the provided reference texts, or to select a tie if both models' outputs seem equally correct (or incorrect).

To combat the issue of positionality bias, in which the judge is biased towards either the first or second model, we provide each comparison twice, once with PROCYON first and once with PROCYON second. We then only use results in which the judge's decision is consistent across both permutations. In the case where the judge selects a tie in one permutation, and a winner in the other, we use the single winner as the result of the comparison. To avoid self-bias from any of the text-only LLMs we compare against, we use a different model for judging, Anthropic's Claude-3.5-Sonnet. The full prompt used to generate the LLM-based evaluations is provided in Supplementary Note 15.

**Limitations.** LLM-as-a-judge is still an experimental methodology for LLM free-text evaluation, and the results should be considered very cautiously. While widespread in evaluation of general question-answering tasks [103], LLM-as-a-judge has not yet been proven in scientific settings. In particular, recent literature has identified that LLM-as-a-judge is not reliable for novel scientific research ideas [104]. In addition, the results of this evaluation may be biased by factors unrelated to the "correctness" of the model's output, including writing style and other latent distributional differences that bias the judge's output. However, we release these results in full transparency of our model's performance, but we acknowledge that this metric alone is not sufficient to judge the performance of PROCYON or the baseline models.

#### 4.7 Domain prioritization for drug targets

In Figure 5a,b, we show how PROCYON can be used to find the binding domain for drugs given the protein targeted by the drug. We extract experimentally validated domain-drug interaction data ("DrugDomain_v1.0_data_PDBs_ECODv291.txt") from [46]. We consider only domains families and proteins in our database. As some proteins have multiple sections of the same domain family, we map the domains to specific domains on the protein by their positions on the protein, as provided by `pdb_start` and `pdb_end` in the data. To account for annotation differences across protein databases, we accept a mapping as long as `pdb_start` is before the start of the domain as annotated in our domain database, while `pdb_end` is after the end. Interacting residues are further compared to obtain the shift, when possible. This processing workflow leaves us with 521 samples (pairs of drugs and domains). The final samples are (drug, protein) pairs with a mapping from the protein to a domain on which the drug binds; thus, each sample contains a set of domains

$(D_1, \ldots, D_i^*, \ldots, D_N)$, where all $D_i$ are domains and $D_i^*$ is the binding domain for the drug.

After these processing steps, we then filtered by a few criteria. First, we filtered for all samples where there were at least 2 Pfam domains that did not bind to the drug for that given sample. Thus, all samples considered contain proteins have at least three domains, including the binding domain. Second, for simplicity, we filter out all instances where multiple domains bind the drug. After both filtering steps, we're left with 35 samples total.

We construct input instructions in the same manner as for drug-target retrieval during training, using identical instruction inputs. However, when we compute the retrieval scores, we score them over the set of domains for the given protein. Please see Methods Sec. 2.3 for more information. For computing results, we calculate the absolute rank (an integer, where 1 is the best) of the binding domain in the prioritized list. The ranking is determined by the similarity score output by the model. We calculate these ranks across all 35 samples in the dataset and plot them in a bar plot in Figure 4b. We compare to ProtST for this task in Supplementary Figure 8. To aid in understanding of the analysis, we also show a few more examples of lists with the correct binding domains, including one where PROCYON misidentifies the binding domain as rank 2 in the list.

We further compare our results to a simple random reference. Importantly, we filter for multi-domain proteins in this experiment, so a random ranking, i.e., a model that does not represent understanding of drug-domain binding properties, is unlikely to produce a similar ranking to that of PROCYON, given the number of domains for each protein. Therefore, the random reference takes the expected rank of the correct binding domain (the middle of the list, i.e., the list size divided by two), and subtracts the actual rank produced by PROCYON from it. Thus, we see that Rank 1 proteins are on average 2.5 ranks above the expected random rank. Full results for this experiment are included in Supplementary Table 30.

#### 4.8 Protein-peptide binding prediction

In this analysis, shown in Figure 4c, we examine the transferability of PROCYON to a novel task and therapeutic modality: protein-peptide binding prediction. Further, we test PROCYON on synthetic peptides, demonstrating that the model generalizes to large distribution shifts at test time. The results for this experiment are shown in Figure 4c.

First, we curated a database of protein-peptide physical interactions from PDBBind. Specifically, we followed the training set evaluation from CAMP framework [49]. This dataset contains 1565 tuples of proteins-peptides, containing the conformations of the complexes, as well as the corresponding binding affinity scores. First, we filtered the original dataset, so that it only consists of pairs of proteins-peptides, since a subset of the samples consists of more than two structure components. This resulted to a total of 1202 protein-peptide samples. Moreover, we included the union of inhibition constant $K_i$, dissociation constant $K_d$, and half-maximal inhibitory concentration $IC50$ values for each protein-peptide complex. It is noteworthy that a subset $(680)$ of peptides in the protein-peptide dataset have their own accession Uniprot ID. In these subset of samples, the interactions were treated as PPI relations. After filtering, this dataset consists of 1097 examples of protein-peptide complexes. The use of this database is notable as PROCYON is not trained explicitly on physical binding data during training. PROCYON is trained on protein interactions from STRING database; however, these interactions and similarities are limited to homology, coexpression, and experimental interaction data, so physical binding data is needed to tune the model for this specific task.

Second, we finetune PROCYON-FULL on this data. We design interaction instruction templates for both QA and retrieval in the same manner as for other interaction datasets during training (see Methods Sec. 3.1 for more information). Thus, PROCYON is tuned on only QA and retrieval with the PDB binding data to produce PROCYON-BIND, which predicts physical protein-peptide binding interactions. PROCYON-BIND is trained for 6.54 epochs for both QA and retrieval tasks on the PDB binding dataset. More details about this model are included alongside hyperparameter reports for other versions of PROCYON (Supplementary Information).

Next, we curate a dataset of interactions between synthetic peptides and angiotensin converting enzyme 2 (ACE2). Our dataset, previously generated by Zhang et al. [50], consists of 58 detected binders for ACE2 and 5072 non-binders taken from a large randomized library of synthetic peptides generated with similar methods gathered by Brown et al. [51]. All peptides are 13 amino acids long due to the library's synthetic construction.

As an initial test of this dataset, we perform testing of the peptides on the original ESM-2 3B model [32]. To test ESM-2's capability to identify these peptides, we test use the proximity of the embedding of each peptide to the embedding of ACE2 as a binding prediction score. ESM-2 achieves a very poor 0.3923 AUROC. This matches results seen from Brown et al. [51], which also found the identification of these ACE2 binders to be challenging for unsupervised machine learning models.

Next, inference is performed to rank peptides for their likelihood of binding to ACE2. Our prompts for both QA and retrieval for this task are identical to that used during training on the PDB

Binding data, including the same in-context examples and the same task definition (see Methods Sec. 3.1). We perform both QA prediction and retrieval prediction for the peptides included in the ACE2-binding set. Importantly, the binding nature of each peptide (i.e., if it is a binder or not) is not given to the model during inference but rather is predicted by the model. For QA scores, we take the difference between the "yes" and "no" token outputs as the singular score of the model; this normalizes the score to a range between [-1,1] where a lower score is considered less likely by the model to be an ACE2 binder and higher is considered more likely to be an ACE2 binder.

For retrieval, our query takes in ACE2 as the input and the retrieval embedding for the prompt is compared to all embeddings of the peptides, thereby calculating the retrieval similarity score comparing the input to all candidate peptides. Each peptide is then ranked by its similarity score, where a higher similarity score corresponds to a prediction of being more likely to be a binder and a lower similarity score is less likely to be a binder. As this procedure follows our typical retrieval inference procedure, we refer the reader to Methods Sec. 2.3 for more details.

We then combine the QA and retrieval results to arrive at the final binding score. The rationale for this comes from the geometry of the latent space whereby there is no observable linear separation between binders and non-binders, thereby hindering the maximum capacity of retrieval in this case. Such lack of separability is most likely due to the challenging nature of this task and that this set of synthetic peptides is indeed out-of-distribution compared to the PDB-binding dataset used to finetune ProCyon-Bind. QA is not limited by this lack of linear separability, but we retain the retrieval scores since retrieval performs very well during training of ProCyon-Bind and thus should recover some signal of the binders. To combine QA and retrieval scores, we first normalize each score to $[0, 1]$ based on min-max normalization; then we take the mean of each score, i.e., $(s_{\text{QA}} + s_{\text{ret}})/2$. This produces the final combined "Predicted Binding Score" from ProCyon-Bind. We then plot the distributions of scores for binders and non-binders beside each other and conduct a Mann-Whitney U test for statistical significance. The final results of the separation gave 0.6480 AUROC for separation of binders and non-binders for the ACE2 dataset while the Mann-Whitney U test showed that ProCyon-Bind achieved a statistically-significant separation of binders and non-binders ($p = 1.04e - 4$).

**Limitations.** In this analysis, we do not claim ProCyon-Bind is a state-of-the-art protein-peptide prediction model. Many models exist for prediction of protein-peptide binding, including AlphaFold3 [1]. Rather, this analysis demonstrates the transferability of ProCyon to new tasks, including those for which it was not trained and on an entirely new modality than its training set, i.e., peptides.

#### 4.9 Retrieving with composite queries across knowledge domains

In this analysis shown in Figure 4d, we examine the abilities of ProCyon to perform composite queries across knowledge domains, performing drug target retrieval that is conditional on the indication of the drug, i.e., the disease that the drug is treating. We gathered descriptions of Nicotine addiction and MDD from StatPearls, focusing on molecular information about the disease, i.e., not clinical diagnosis information. On August 08, 2024, we retrieved the StatPearls description for Depression from the "Pathophysiology" section on the webpage entry [105]. We used the depression entry [59] instead of MDD because the MDD description [105] does not provide explicitly a "Pathophysiology" section. In addition, since MDD corresponds to a subtype of depression [41] with a relatively strong biological component, the "Pathophysiology" section of depression was almost exclusively focused on MDD. On August 8, 2024, we retrieved the StatPearls description for Nicotine addiction [58]. The exact descriptions used in the forward pass to the model are given in Supplementary Note 17.

#### 4.10 Phenotype generation for poorly annotated proteins

This section addresses the generation of phenotypes for AKNAD1 and other proteins in Figure 5a,b. To generate these outputs, we use a simple phenotype generation query for ProCyon that copies the structure of the prompt used for UniProt during training. In the input prompt, we insert the proteins AKNAD1 (UniProt: Q5T1N1), LYRM2 (Q9NU23), ANKDD1A (Q495B1), and UQCC4 (Q4G0I0). We use ProCyon-Full for this prediction. AKNAD1 was chosen due to the fact that it is not in our training data, but external evidence of its function exists in the Human Protein Atlas (HPA). Thus, this is a zero-shot prediction for ProCyon on a protein that has been poorly annotated in literature. Other proteins were chosen from proteins without annotations at the time our UniProt data curation (December 2022)

We use diverse beam search to generate the phenotype, and we generate 10 generated phenotypes for this protein. After generating these descriptions, we perform QA filtering (see Methods Sec. 2.3) to derive the top-3 captions for this protein. This gives us the three descriptions seen under label "Generated Phenotype".

Next, we validate the generated phenotypes for this protein with immunofluorescent and expression evidence through the Human Protein Atlas [8] (Human Protein Atlas proteinatlas.org). For the top description, we find immunofluorescence evidence supporting that this protein is localized to the cytoskeleton. Specifically, immunofluorescent staining of AKNAD1 (antibody HPA030270, https://www.proteinatlas.org/ENSG00000130590-AKNAD1/antibody) in cell line HAP1, which is a "near-haploid human cell line derived from KBM7, a human myeloid leukemia cell line developed from a 39-year-old male patient in the blastic phase of chronic myeloid leukemia" has confirmed its localization to cytosol, microtubules, and cytokinetic bridge.

For the second description, we find two lines of gene expression evidence supporting that the protein is overexpressed in spermatids, leading to a likely conclusion of the protein being involved in spermatogenesis. In a meta-analysis of single-cell RNA sequencing literature and databases conducted by HPA [8], a group enrichment (nTPM at least four times any other cell type) of the expression of AKNAD1 is observed in both early spermatids (90.9 nTPM) and late spermatids (29.6 nTPM). In addition, inferred from RNA-seq of unfractionated tissue samples [106], very high enrichment of AKNAD1 expression in late and early spermatids were observed (mean correlation=0.696, 0.695, respectively, with 3 reference transcripts selected to represent the two cell types, specifically CEP55, KPNA5, PBK (late spermatids); PRM1, PRM2, TNP1, respectively, data extracted through HPA [8] (Human Protein Atlas proteinatlas.org).

#### 4.11 Systematic prediction of functions of poorly characterized proteins

In this analysis (Figure 5c,d), we focus on predicting functions for poorly characterized proteins, defined by an annotation score no higher than 3 on UniProt at its curation time in ProCyon-Instruct (Dec 2022) and no annotation across all protein annotation sources (GO, Reactome, EC, DrugBank, GtoP, DisGeNet, OMIM) in ProCyon-Instruct. We also ensure that:

- The proteins have transcript- or protein-level existence evidence.
- Their associated genes induce strong transcriptional phenotypes as defined by having at least 50 differentially expressed genes at a significance of $p$-value $< 0.05$ by Anderson-Darling test following Benjamini-Hochberg correction in [61] (first criterion from the original paper, data courtesy of authors).
- The proteins do not have identified domains, or their identified domains are either uncharacterized (domains of unknown function (DUF) or uncharacterized protein families (UPF)), or in the test set.

- The proteins were not discussed in [61].

These stringent filters ensure the proteins we investigate have limited evidence in curated databases, but can be validated with perturbation data. Nine proteins (see Figure 5d) are left for analysis. We then query PROCYON (PROCYON-Split) for functions that are in our database and calculate the retrieval ranks of these poorly characterized proteins. To validate such predictions, for each poorly characterized protein, we perform gene set over-representation analysis on the differentially expressed genes upon the corresponding knockdown for established gene sets "GO_Biological_Process_2023" and "GO_Molecular_Function_2023" from Enrichr [62] using GSEApy [107], with all genes measured in [61] as background. Functions that were significantly enriched (adjusted $p$-value $< 0.05$) with notable enrichment (odds ratio $\geq 2$ and gene overlap $\geq 3$) were deemed identified from perturbation data.

#### 4.12 Analyzing poorly-characterized proteins for Parkinson's Disease

This section encompasses the analysis described in Figure 5e and Extended Data Figure 9, where we study the pathways predicted by PROCYON for poorly characterized proteins that are hypothesized to have association in Parkinson's disease. Analysis of these experiments is described in Results.

**Gathering the PD-uncharacterized list.** We seek to study the ability of PROCYON to uncover phenotypes of proteins that are poorly functionally characterized in literature. To get this list, we gather all proteins in OpenTargets that are hypothesized to be associated to Parkinson's disease (PD) [66] (Accessed on the 12th August 2024). We filter the list at those above 0.1 association score threshold. We then cross-reference this list with the proteins in our database, and we gather the proteins in the list from OpenTargets that do not appear in any of our training databases. This creates the PD-uncharacterized list, a list of 36 proteins with a link to Parkinson's disease that are considered **zero-shot examples** for PROCYON since it was not trained on any phenotypes for these given proteins. This full list is given in Supplementary Table 31.

**Generation of descriptions with PROCYON.** We follow the standard pipeline to generate descriptions of the proteins in PD-uncharacterized. We use prompts in the style of GO biological process to generate descriptions of the pathways in which each protein might be involved. We use QA filtering with a threshold of 0.6 to get the final list of descriptions for each protein.

**Automatic clustering.** The automatic clustering is performed by grouping together similar descriptions that have a high n-gram overlap similarity. First, we take ROUGE-L scores for every pair of generated descriptions $\{(d_i, d_j) | d_i, d_j \in \mathcal{D}\}$, resulting in a $(|\mathcal{D}|, |\mathcal{D}|)$ distance matrix. This distance matrix is then converting into a binary adjacency matrix by taking all distances greater than 0.95 (which are considered to be "copies" by ROUGE-L) as edges between descriptions. A connected components algorithm is then performed on the resulting graph to find all connected components, i.e., groups of descriptions that are similar to each other. This results in 9 clusters, with an minimum membership of 3 and a maximum membership of 7. These clusters are then used to collapse down the number of descriptions for each of the proteins in the PD uncharacterized set.

**Manual clustering and curation of pathways.** The protein descriptions that are not assigned to a cluster through the automatic approach are manually evaluated by two experts. The descriptions are annotated independently by two experts and assigned a cluster label (i.e., a pathway) according to currently available literature (see Supplementary Note 20). The two independent annotations are examined for consensus, and a unique label is then derived. The same approach is followed to assign pathway labels to the automatically derived cluster, where the group descriptions rather than individual protein descriptions are used. Finally, the proteins with the same manually assigned pathway annotation are grouped together.

**Enrichment testing.** To test quantitatively whether the pathways generated by PROCYON are implicated in PD, we test pathway enrichment among a gene list of well-known PD-associated genes ("PD-Associated") to two control groups: one of neurologically-expressed genes ("Neuro control") and one of genes expressed outside of neurological tissue ("General control") . Exact details on these lists is included in Supplementary Note 19.

We then use these lists to compute the enrichment of each pathway derived from PD-uncharacterized within the lists. First, we calculate a proportion expressing the coverage of each GO term by the proteins in the list. The proportion is calculated as the number of proteins in the given protein list $\mathcal{P}$ that are associated to the cluster's assigned GO term $\mathcal{G}_i$ over the size of that GO term:

$$\text{Proportion}(\mathcal{G}_i, \mathcal{P}) = \frac{\sum_{p_j \in \mathcal{P}} \mathbf{1}_{\mathcal{G}_i}(p_j)}{|\mathcal{G}_i|}; \mathbf{1}_{\mathcal{G}_i}(p_j) = \begin{cases} 1 & \text{if } p_j \in \mathcal{G}_i \\ 0 & \text{else.} \end{cases} \tag{7}$$

By dividing by the size of the GO term as opposed to the size of the control list, this formula considers the "specificity" of the pathway, i.e., how many proteins within the pathway are found in these given lists. We can then directly compare the number of proteins in a given list to another list by subtraction, i.e., Proportion$(\mathcal{G}_i, \mathcal{P}_1)$ − Proportion$(\mathcal{G}_i, \mathcal{P}_2)$. We calculate this difference for PD-associated vs. the neuro and general lists, respectively. We then show these differences of proportions as bars in Figure 6b.

To balance the statistical comparison we perform downsampling of both the neuro and general control lists during computation of the proportions. This is built off the intuition that a large randomly-gathered list of genes would have a high likelihood of including the associated proteins in the GO term since our proportion calculation (Equation 7) does not account for size of the list $\mathcal{P}$. Thus, we randomly sample each control list to the size of the PD-associated list (962) 30 separate times, then we take the median proportion from those 30 separate trials.

To compare statistical significance of the proportions, we perform a one-sided proportion z-test where the alternative hypothesis is that the proportion in the PD-associated list is higher than in the control list. We show the results of this significance test in Figure 5e as well as in Supplementary Tables 32 and 33. In Figure 5e, we highlight the bar in the barplot as green if the difference in proportions is indeed significant.

**Describing poorly-annotated proteins identified through alpha-synuclein screens.** Next, we examine a list of proteins that were discovered to be involved with the molecular pathways leading to the toxic effect of $\alpha$-synuclein. We first gathered data from a study that ran $\alpha$-synuclein toxicity screens on yeast and then transferred the discovered genes based on homologous proteins in humans. Specifically we gathered the list of proteins from Supplementary table S9 in Khurana et al. [68]. Then, we filtered for proteins that were not present in our training data, i.e., proteins on which our model had seen no linkage to phenotypes. This left us with only four proteins, ANKDD1A (UniProt: Q495B1), SVOP (UniProt: Q8N4V2), TCEAL5 (UniProt: Q5H9L2), and ZNF654 (UniProt: Q8IZM8). We then generated descriptions in both the style of the UniProt database (i.e., general functions), and in the style of GO biological process. We selected a few descriptions to show in Extended Data Figure 10. We show the protein structures as well as generated descriptions below the structure and sequence of each protein. Protein structures are gathered from AlphaFoldDB [108], and we include the pLDDT scores (i.e., confidence of the structure at a per-residue granularity level) to illustrate the poor annotation of the proteins from a structure perspective.

#### 4.13 Interrogating multiple sclerosis targets on an experimentally-derived dataset

In Figure 6, we show the PROCYON-driven analysis of a novel experimentally-derived dataset. In this section, we describe the generation of this dataset as well as how we used PROCYON to interrogate genes found in the experimental procedure.

**Dataset generation and neuropathological assessment.** Clinically and neuropathologically confirmed MS brain donors were recruited through the UK MS Society Tissue Bank (MSSTB) at Imperial College, London. Autopsy procedures were carried out after obtaining fully informed consent and under ethical approval granted by the National Research Ethics Committee (08/MRE09/31). Dissection of Superior Frontal Gyrus (SFG) blocks was performed for all subjects following a standardized protocol, as previously described [109]. This region was selected due to its well-established significance in MS pathology [110, 111]. Cases were processed for luxol fast blue histology and anti-myelin oligodendrocyte glycoprotein immunostaining to characterise grey and white matter and areas of demyelination. Sections were immunostained for human leukocyte antigens using a mouse anti-HLA (DP, DQ, DR; cr3/43; Dako, Glostrup, Denmark) and detected with an anti-mouse peroxidase linked secondary antibody (Immpress HRP, Vector Labs, UK), with diaminobenzidine as the chromogen. The percentage area of HLA-D+ immunostaining was calculated from non-lesion white and grey matter (minimum area =5mm2) per donor block using the positive pixel classifier tool in QuPath (v0.5) [112] as previously described [113]. Anti HuC/D (clone 16A11) immunostaining revealed post-mitotic neurons (see Figure 6). RNA was extracted from the gray matter of 225 SFG blocks with the Qiagen RNAeasy Tissue Lipid Mini kit, following the manufacturer's protocol. Library preparation was conducted using the Lexogen Quantseq 3' mRNA FWD kit with dual indexing, and sequencing was performed on the Nextseq2000. The RNA-seq workflow from the nf-core framework (v3.3) was performed using Nextflow (v21.04.0) [114]. Raw sequencing reads in fastq format were aligned to the GRCm38 genome assembly using STAR (v2.7.6a) in conjunction with the corresponding Ensembl transcript annotations [115]. The resulting BAM files were sorted by chromosomal coordinates using samtools (v1.12). Finally, RSEM (v1.3.1) was employed to compute gene-level read count estimates and to derive TPM values [116]. After these steps, 222 samples were retained for downstream analysis. Outlier detection was performed on the first principal component (PC1) using an IQR-based method. Specifically, samples with PC1 values falling beyond 1.4 times the interquartile range

(IQR) from the 25th and 75th percentiles were considered outliers. This threshold was selected based on a systematic evaluation of various thresholds, where the 1.4 multiplier provided a rigorous criterion that minimized undue sample loss while clearly identifying samples whose inclusion distorted the overall data structure. The identified 19 outlier samples demonstrated extreme deviation from the main distribution of the data, and their removal resulted in a more coherent clustering of the remaining samples. These 19 outlier samples were identified corresponding to samples with RIN values below 4 or unique read counts below 1.5M and were subsequently removed from further analysis. Additionally, 7 samples were excluded due to incomplete metadata.

**Differential expression analysis.** Differential expression analysis was performed using the edgeR package, adjusting for sex and RIN score. The independent variables included the log transformed HLA staining degree in grey (n = 174) and white matter (n = 174), presence of active demyelinating regions (n = 196), percentage of SFG white matter lesions (n = 183) and presence of meningeal and perivascular inflammation (n = 196). The workflow involved filtering out lowly expressed genes, normalizing library sizes, estimating dispersion, and fitting a generalized linear model with a quasi-likelihood F-test. Genes were considered significantly differentially expressed when the false discovery rate (FDR) was below 0.05. The volcano plot of DEGs for HLA+ in grey matter is shown in Figure 6g and the rest are shown in Supplementary Figure 12.

**Multivariable validation of ProCyon against experimental results.** Next, we validate the retrieval capabilities of ProCyon against prompts describing neuropathological MS hallmarks. We test five different MS hallmarks: HLA+ in grey matter tissues in the SFG, HLA+ is white matter tissues in the SFG, presence of active demyelinating lesions, percentage of white matter lesions in the SFG, and presence of meningeal and perivascular inflammation. We then wrote five prompts describing each of these hallmarks; these prompts were reviewed by a neuropathologist prior to running them through ProCyon. We then used these prompts as inputs to ProCyon for retrieval (OMIM-based instructions) and calculated the percentile score for each DEG with respect to the given variable. Exact prompts are given in Supplementary Table 9.

**Filtering DEGs for HLA+ in Grey matter.** We focus our analysis on HLA+ expression in grey matter, both due to its relative underrepresentation in MS research and because our RNA-seq data were derived from grey matter, enabling more reliable detection of DEGs with reduced confounding from tissue heterogeneity [117, 118]. Of the 23 identified DEGs for HLA+ in grey matter, we looked into the properties of each of these to determine which were the most ”unexpected”, i.e., which genes had not been identified previously as MS-related or involved in common MS pathways (i.e., inflammation and immune response). This filtration process is depicted in Supplementary Figure 15, along with the exact genes filtered at each step. Our first step of filtering involved removing all genes known to be MS-related; MS relation was defined as appearing in the registered list of targets in the OpenTargets [66] database at a ”global confidence score” above 0. In addition, since we focused on the Grey matter, we extracted GTEx TPM counts data from brain regions that are predominantly Grey matter - nucleus accumbens, brain caudate, anterior cingulate cortex, brain cortex, putamen, amygdala, hippocampus, frontal cortex. A threshold of 10 on the TPM count was applied, and genes with a TPM count $> 10$ in at least across 80% of the samples in any of those brain regions were extracted as Grey Matter genes. These genes were intersected with the MS genes to identify the subset of MS genes that are expressed in the Grey Matter, which were then used as “MS-related” genes for filtering. This step removed 11 genes. Second, we removed all genes that are directly associated to inflammatory and immune responses. The rationale at this step is centered around the mechanism of HLA+ staining: HLA molecules are immune and inflammatory biomarkers, and thus many genes involved in inflammatory and immune processes might get identified as being differentially-expressed, but are not directly involved in MS. Practically, this was done by removing all genes annotated to gene ontology terms immune response (GO:0006955) and inflammatory response (GO:0006954). This step removed an additional 3 genes. To account for the GO hierarchy, we also propagated relations such that if a gene was associated to child terms of immune or inflammatory response, it was also considered as annotated to those terms themselves. This step removed 1 gene. After removing this final gene, we were left with 8 ’unexpected’ genes which did not have a direct explanation for their involvement in MS nor inflammation and immune response.

**Measuring MS relevance versus strong control disease sets.** We next conducted a disease enrichment analysis to identify whether PROCYON associated these ’unexpected’ genes as being more closely related to MS or other diseases with different degrees of similarity to MS. We considered 4 groups of control diseases: central nervous system (CNS) inflammatory diseases, known inflammatory and immune-related diseases with shared pathways to MS, known inflammatory or immune-related diseases with no shared MS pathways, and finally diseases which are not inflammatory or immune-related and share no pathways with MS. The full list of diseases for each group is available in the Supplementary Table 9. We extracted disease descriptions from OMIM for all categories except CNS, where the specificity of included conditions required sourcing descriptions directly from the primary literature (full references provided in Supplementary Note 28). These descriptions were input into ProCyon for protein retrieval with prompts similar to those used to train on OMIM-based instructions. For each "unexpected" gene, we recorded its ranking percentile across diseases and ranked the diseases accordingly, with higher percentiles (closer to 100) indicating stronger enrichment for the corresponding protein. To benchmark these rankings, we also retrieved known gene–disease associations from OpenTargets for each gene and its respective control diseases, allowing us to assess whether ProCyon ranked MS even higher than well-established associations. Details on these associations are available in the Supplementary Table 34. To assess statistical significance, we performed a permutation-style analysis using three control gene sets: a) non-DEG, non-inflammatory, non-immune genes; b) a subset of (a) further excluding MS-associated genes; and c) a subset of (a) limited to known MS-associated genes. Sets (b) and (c) together comprise the full set (a). Gene classifications for inflammation, immunity, and MS association followed the criteria described above.

For each control gene, we repeated the same disease enrichment analysis as applied to the "unexpected" genes. To compute a p-value for each unexpected gene $i$, we calculated the proportion of control genes with equal or higher MS ranking:

$$p_i = \frac{\text{Number of control proteins ranked } \geq r_i}{\text{Total number of control proteins}}$$

This analysis was conducted independently for each control set to ensure robustness of the enrichment signal across different baselines. These analysis resulted in 3 genes being significantly enriched relative to baselines, which were used for further analysis.

**Identify potential pathways related to MS.** To identify potential mechanisms linking the three "unexpected" genes to MS, we perform a Gene Ontology (GO) enrichment analysis. We input each GO term into ProCyon for protein retrieval and record the ranking percentile of each gene across all GO terms. For each gene, we select the top 15 GO terms in which it ranks highest relative to other proteins. For each of these top GO terms, we calculate an MS enrichment score as:

$$\frac{\text{\# of MS proteins in pathway}}{\text{total proteins in pathway}}$$

Where MS-associated proteins are defined using the Open Targets MS gene list. A neurologist conducted a qualitative assessment of the top 15 ranked pathways for each of the three genes.
